# Cuticle and cuticular sensilla in Agnostina

**DOI:** 10.1101/2022.03.25.485613

**Authors:** Elena B. Naimark, Stanislav Yu. Chaika

**Affiliations:** Paleontological Institute RAS, Profsouysnaya, 123; Lomonosov State University, Biology Department, Vorobievy Gory 1, Moscow, Russia

**Keywords:** arthropods, Cambrian, Agnostina, cuticle, exoskeleton, sensilla.

## Abstract

The microstructure of the cuticle of several Agnostina species is described. For the first time, it has been shown that their cuticle consists of three layers: thick outer and transitional layers and a thin inner layer. The layers are made up of stacks of laminae: the laminae in the outer layer are thicker than in the transitional and inner ones. The evolution of the agnostoid cuticle proceeded, apparently, in the direction of decreasing the thickness of the inner layer which in contrast to all other arthropods, is thinner than the outer layer. Apparently, the inner layer was mineralized to a lesser extent than the outer one. In the structure of the cuticle, Agnostina are close to horseshoe crabs and trilobites. On the outer and inner layers, a set of sensilla was revealed; they can be attributed to trichoid, campaniform, and digitiform sensilla (with mechanoreceptor function) and celoconic sensilla (with chemoreceptor function). Also, unique sensorial ensembles were found on cephalons and pygidia; the function of the former is probably mechanosensorial, and the purpose of the latter cannot be assumed on this material.

---

*Agnostus pisiformis* (Wallerius, 1818) and other Agnostina (Euarthropoda; hereinafter agnostids) (Müller and Walossek 1987; Moysiuk, Caron 2019) miniature blind inhabitants of open waters, were quite common in the middle and upper Cambrian throughout the world and are an important stratigraphic group for a given time interval. Agnostids belong to arthropods, a group for which the exoskeleton is one of the primary anatomical features. The structure of the shell in arthropods varies; it reflects both the phylogenetic relationship of groups and adaptation to living conditions. However, despite the fact that agnostids have been studied for more than two centuries and there is no shortage of material on them, surprisingly little is known about the structure of their shell.

The agnostoid exoskeleton was studied mainly on examples of *A. pisiformis* and *Homagnostus obesus* (Belt, 1867) supplemented with single specimens of Peronopsis and Ptychagnostus (Wilmot 1988, 1990). It has been considered to be built up from one thin (5-15 μm) cuticular layer. This layer had been assumed to be lamellar or prismatic (Wilmot, 1990) or irregularly mosaic (Tegleir, Towe 1975). On the material of *H. obesus*, Wilmot indicated the presence some cavities or pits on the visceral side of this single cuticular layer (Wilmot 1988: pl. 6.1, fig. F). The functions of these depressions remained unclear.

The seminal work on the phosphatized soft tissues of *A. pisiformis* also concerns the details of the structure of their exoskeleton. Although the authors could not clearly determine the number of cuticular layers (one or more), they described a surface polygonal pattern, openings of different sizes, and some cuticular structures, which, with some doubt, they assigned to sensilla (Müller and Walossek 1987).

As far as we know, there are no more data on the structure of the agnostoid carapace. We present here the SEM images and a description of cuticular architecture and microstructures in several representatives of agnostids. Studied specimens are few however they allowed us to carry out careful morphological descriptions of cuticular arrangement and compared it with other groups of Arthropoda.

In contrast to previous views, the cuticle of the agnostids now appears to have a rather complex arrangement. In our accompanying paper we described in details the cuticle of one agnostoid species (Naimark, Chaika, 2022), and here we extend the description on the material of other species from different agnostoid families. The cuticles vary across families, however its common features indicated a relationship with trilobites, eurypterids, and xiphosurians.

We also revealed an unexpectedly diverse suit of sensilla, pores, and sensorial fields in the cephalic and pygidial shields. Given this diversity, agnostids are now appearing to be not blind insensate creatures, but animals equipped with special detectors that were tuned to get signals from the outside environment.

## MATERIAL AND METHODS

The material came from the Chekurovka section, in east Siberia (Fig. 1). The specimens were chosen from the very rich collection, which comprises the upper Cambrian strata (Korovnikov, Novozhilova 2012; Lazarenko *et al*. 2011).

**Fig. 1.**
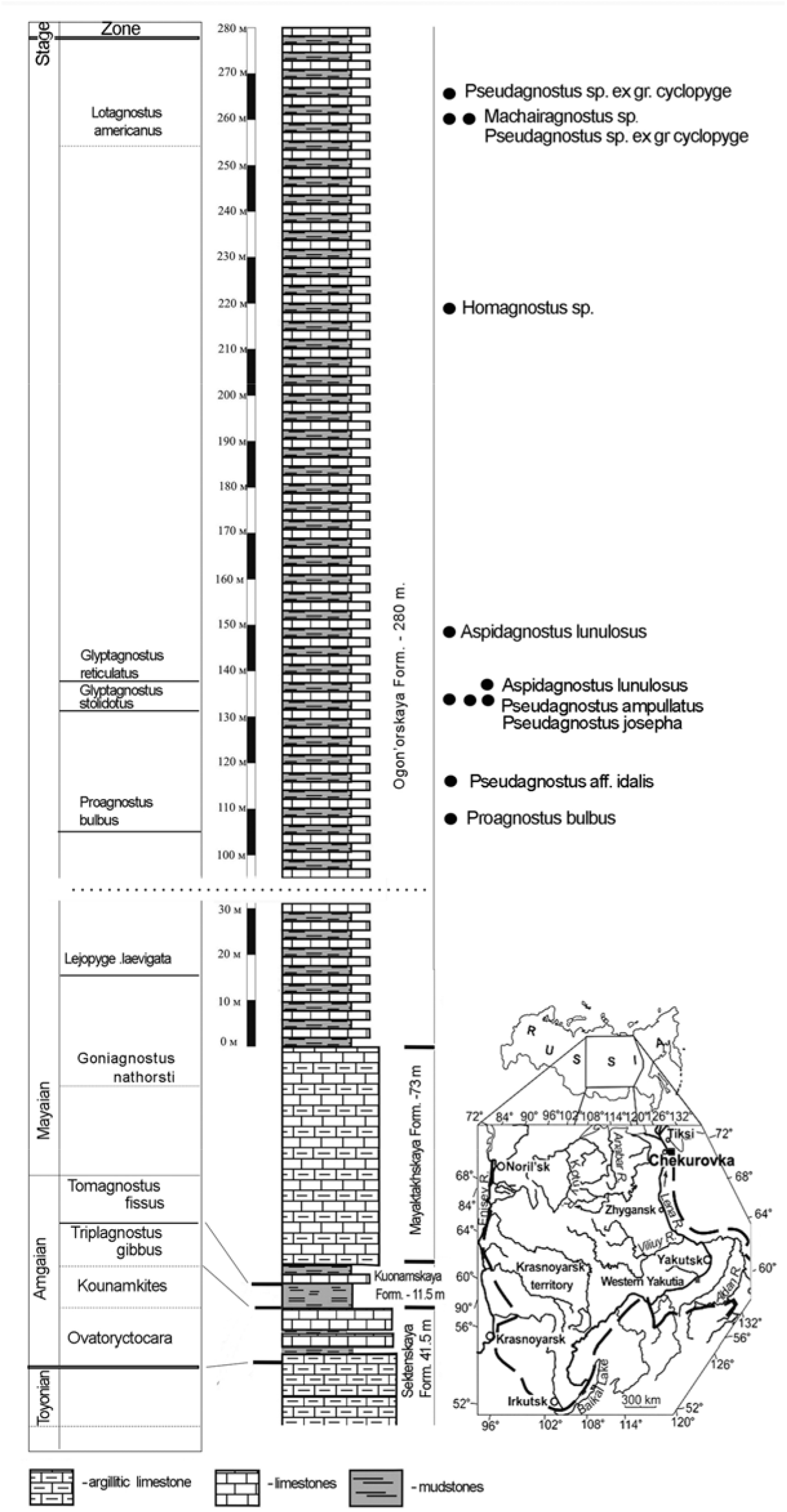
Сhekurovka section in the Ogon’or Fm., northeast of Siberia, the positions of the samples within the section are shown. This unpublished scheme was created by I.V.Korovnikov and E.B.Naimark based on Lazarenko *et al*. 2011 and Korovnikov, Novozhilova 2012.

Our choice of specimens was random; it was dictated by the presence of any interesting details on the exoskeletons visible under a light microscope with a magnesium coating. The chosen specimens belong to the families Agnostidae, Ammagnostidae, Aspidagnostidae, and Pseudagnostidae and included cephalons and pygidia. We defined the species using the entire rich collections from the corresponding beds.

The chosen specimens were investigated under SEM with a palladium/gold coating. For most specimens point elemental analyses were provided (with Zeiss EVO-50 and associated energy dispersive X-ray spectroscope INCA Oxford 350 at 15 kEv).

The specimens are stored in the Paleontological Museum in Moscow, collection N5862.

## RESULTS

The cuticle of the agnostids is subdivided into three layers. What we call layers are those cuticular divisions along which the cuticle splits most often. Laminae are thin horizontal or oblique units in the layers. No other morphological or biochemical meaning for layers and laminae is implied in this work.

When identifying cuticular microstructures, it is important not to confuse them with crystal defects or post-mortem changes, including microdrillings. Therefore, we focused on those elements that occurred more than once and had a regular shape. Below we give brief descriptions of each specimen studied (Fig. 2-10); the captions in the figures also provide some interpretations to the most interesting elements displayed in the images. The results of point elemental analyses are summarized in Table 1 and Naimark, Chaika, 2022b.

**Fig. 2.**
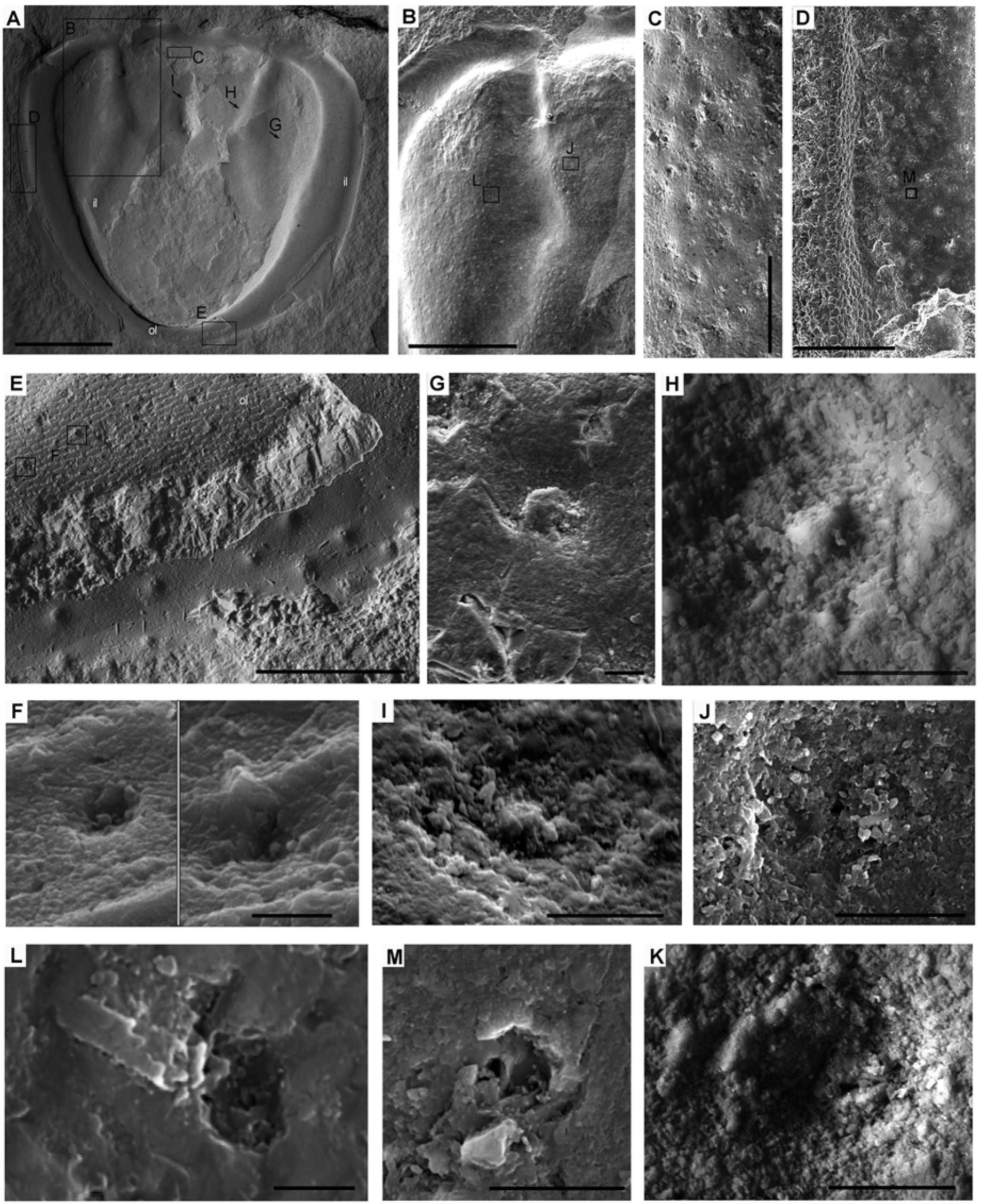
Pygidium *Pseudagnostus aff. idalis* PIN 5862/10. A, general aspect; here and below, framed fragments and arrowed locations that are indicated by small letters are enlarged in the corresponded panels. B, the fragment of the left part of the acrolobe; a small piece of the outer cuticular layer of the border is visible on the left corner (black arrow); the axis and acrolobe bears multiple peg pits visible as shallow indentations. C, the enlarged fragment of the upper part of the axis (turned 90° clockwise from A); multiple pits are visible, some of them retain central pegs. D, enlarged fragment of the edge of the left part of the border; outermost prismatic sublayer of the outer layer is visible. E, fragment of the border furrow and border; a thick piece of the outer layer with surface pentagons is visible; the layer below this piece is the inner layer with multiple regular peg pits. F, small indentations on the surface of the outer layer. G–K, examples of the peg pits variably performed on the inner layer: G, typical appearance of the peg pit, where the peg has a smoothed whitish top; H, some kind of meshed structure is visible on the bottom of the pit. I, the top of the peg is covered by some smoothed substance. J – the central peg is absent, and the tiny opening is visible in its place; K – an opening surrounded by a convex collar. L, a complex structure on the inner layer: a round indentation with a tiny hole containing a rod; a piece of a tube-like shape is next to the ressess; this is possibly the remains of a glandular channel. M, an inner layer: a pore surrounded by a cone-like opening; the remains of the immediate upper cuticular unit is visible in the right; we interpreted it as remains of a trichoid element. Scale bars: 2 mm (A); 1 mm (B); 100 μm (C); 200 μm (D, E); 5 μm (F); 10 μm (G–K, M); 2 μm (L).

**Table 1.** Point elemental analyses of the agnostoid exoskeletons and in the surrounding rock (in atomic %; K, Na are ommited). Values for outer (O), transitional (Trans) inner (I) layers and rock are given separately. All values are averaged across 3-15 measurement; for values at each point see Data repository: Naimark, Chaika, 2022b.

| species | place | C | Mg | Ca | Al | Si | Fe |
| --- | --- | --- | --- | --- | --- | --- | --- |
| *Pseudagnostus* aff. *idalis* | rock | 27,05 | 0,88 | 5,82 | 0,4 | 1,48 | 0 |
|  | O | 28,28 | 0,45 | 5,01 | 0,14 | 1,2 | 0 |
|  | I | 26,16 | 0,33 | 9,98 | 0,12 | 0,2 | 0 |
| *Pseudagnostus* ex gr. *cyclopyge* 5862/1 | rock | 24,50 | 0,61 | 11,65 | 0,18 | 0,45 | 0,09 |
|  | O | 24,82 | 0,58 | 11,44 | 0,13 | 0,36 | 0,05 |
|  | Trans. | 15,02 | 0,30 | 26,86 | 0,08 | 0,14 | 0,00 |
|  | I | 19,44 | 0,39 | 20,06 | 0,09 | 0,17 | 0,01 |
| *Pseudagnostus* ex gr. *cyclopyge* 5862/2 | rock | 23,74 | 0,92 | 10,79 | 0,52 | 1,24 | 0,16 |
|  | O | 23,855 | 0,125 | 14,08 | 0 | 0 | 0 |
|  | I | 26,88 | 1,47 | 8,12 | 0 | 0,03 | 0 |
| *Pseudagnostus josepha* (cephalon) | rock | 19,94 | 0,49 | 13,98 | 0,67 | 2,72 | 0,7 |
|  | O | 19,94 | 0,36 | 19,49 | 0,06 | 0,12 | 0 |
|  | I | 22,48 | 0,31 | 13,01 | 0,65 | 1,33 | 0,14 |
| Machairagnostus sp | rock | 24,20 | 0,67 | 10,38 | 0,49 | 1,26 | 0,15 |
|  | O | 25,63 | 0,25 | 11,12 | 0,06 | 0,07 | 0,02 |
|  | I | 25,26 | 0,29 | 9,95 | 0,38 | 0,86 | 0,12 |
| *Aspidagnostus lunulosus* | rock | 26,66 | 0,54 | 3,25 | 0,92 | 3,06 | 0,25 |
|  | O | 25,01 | 0,37 | 10,76 | 0,34 | 0,5 | 0,06 |
|  | I | 23,99 | 0,58 | 7,82 | 0,94 | 2,61 | 0,31 |
| *Homagnostus sp.* | rock | 35,46 | 0,47 | 8,54 | 0,38 | 1,58 | 0 |
|  | O | 33 | 0,14 | 14,12 | 0,55 | 1,32 | 0 |
|  | I | 30,02 | 0,45 | 21,39 | 0,46 | 0,73 | 0,18 |
| *Proagnostus bulbosus* | rock | 26,07 | 0,16 | 6,18 | 0,59 | 2,02 | 0,79 |
|  | O | 26,72 | 0,25 | 7,08 | 0,39 | 1,23 | 0,26 |
|  | I | 26,00 | 0,20 | 6,89 | 0,55 | 1,94 | 0,31 |

### Pseudagnostus aff. idalis (Opik, 1967) (Fig. 2)

The exoskeleton of this large pygidium (Fig. 2A) displays two layers. The outer layer is represented by small pieces in the rear of the border furrow and along the left edge of the border (Figs. 2A, B, D). These pieces bear a polygonal pattern on the surface. Polygons of around 15 μm are formed by ridges of crystals sized 0.5-0.8 μm. The thickness of the outer layer at the border furrow it is 100 μm; and on the left dublura is around 50 μm (Figs. 2D)

On this upper layer, multiple indentations are present (Fig. 2E, F).

In this specimen, the most of the cuticular surface is represented by the inner layer. Instead of polygons, it is covered by shallow pits (Fig. 2B-K). The axial and border furrows tend to lack these pits, or the pits are rare. These pits have approximately the same diameter (*c.* 15-17 μm). Most pits are smoothed out and only minor depressions remain from them; yet some of them contain a tiny cone (or peg) in the center (Figs 2G – I). If the central cone is absent a small opening is present in its place (Figs 2I, J, K). Complementary to the pits, other structures were found in the inner layer. For example, there is a cone-like opening (Fig. 2M) and a deep indentation with a tiny opening with a rod inside and a tubular element next to this indentation (Fig. 2L).

The elemental contents of the outer and inner layers and of the pits with the central cones (Table 1) showed the same composition with a prevalence of Ca with a small addition of Mg; a comparatively high amount of Si and Al is also present. The host matrix has considerably less Ca and Mg; Si and Al dominated in it. Thus, the inner layer can be differentiated from the host matrix by its elemental content.

### Pseudagnostus sp. ex.gr. P. cyclopyge (Figs 3-6)

This species is illustrated with two large cephalons from two horizons (Fig. 1). On these cephalons, all cuticular structures are represented better than on other specimens in hand. These specimens were described in detail in the accompany paper (Naimark, Chaika, 2022) together with other cephala and pygidia of this species. We repeat here the most important pictures illustrating the morphology of the agnostoid cuticle, and supplementing them with some additional ones which seemed to us informative. To avoid repetitions in the text we only focused on the main cuticular characteristics and the features that were absent in our descriptive paper.

On the cephalon 5862/1 (Figs 3A, B), three layers are variously displayed in different parts of the carapace (Figs 3B, D, L; 4A, C). On the second cephalon N 5862/2 (Figs 4, 5) only two layers can be traced.

**Fig. 3.**
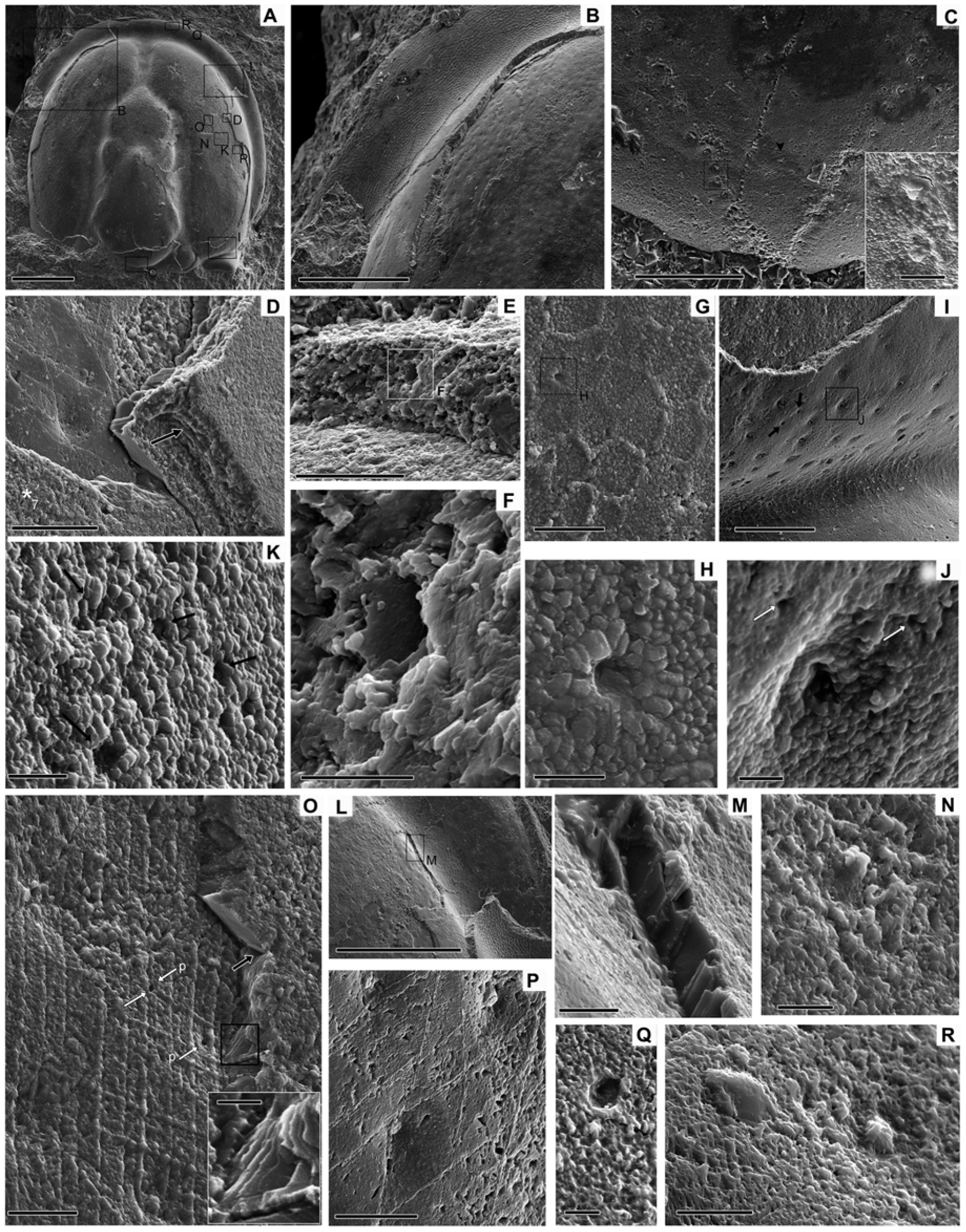
*Pseudagnostus sp. ex gr. P. cyclopyge* PIN 5862/1. A, general aspect. B, the fragment of the acrolobe; subunits of the outer layer are visible. The outermost subunit (the thinnest one) bears the polygonal pattern. The surface of the inner layer covered by shallow pits is visible on a major part of the fragment from the bottom right corner. C, V-shaped groove on the glabellar culmination on the outer layer. In its converged part, the branches of the groove look continuous, however in the diverged part the groove consists of the row of pits with some bulbous bodies (see two of them in the inset). An elongate tubercle centered between the branches (black arrowhead) is discernible (see it in Fig. 12N). D, the fragment shows three layers: the outer layer consisted of the relatively thick lamellae (black arrow), the transitional layer superimposed the inner layer with peg pits. The inner layer is covered by a rhombic mesh. E, the horizontal pore in the outer layer. F, multiple tiny pore canals penetrate very thin wall of the horizontal pore. G, the fragment of the ridged polygons with three rimmed pores; a crystalline texture of this subunit is well pronounced. H, a rimmed pore. I, the fragment of the right acrolobe and the adjusted pleural furrow showing traces of the polygons in the pleural furrow and the pitted surface of the underlaying principal part of the outer layer; small openings are also visible on this surface (black arrows). J, a rimmed large pore and small pores in its vicinity (white arrows). K, a transitional layer with multiple pores (black arrows). L, a fragment where the outer layer with the polygonal surface is clearly discernible as well as the transitional layer; a patch of the inner layer with pits is revealed under it. M, vertical laminae of the transitional layer. N, traces of a peg pit on the surface of the transitional layer; the top of the peg is accentuated by some whitish material while the pit is almost smoothed. O, the transitional layer superimposed over the inner layer; the former consists of stacks of thin lamellae, vertical and oblique to the surface (see the framed fragment in the inset), their junction is shown by the black arrow; the inner layer is displayed here as a rhombic mesh with peg pits, two small pores are visible on this fragment (white arrows). P, the fragment of the inner layer with peg pits; rhombic mesh is clearly visible. Q, a funnel pore in the inner layer of the border. R, a peg pit and an elongate tubercle on the inner layer of the border; note the smooth surface of the tubercle. Scale bars: 1 mm (A); 500 μm (B, L); 100 μm (C; inset 10 μm), 100 μm (I); 50 μm (D, E); 10 μm (F, M, N, R); 20 μm (G, P); 5 μm (H, K); 4 μm (J, Q); 20 μm (O, inset, 5 μm).

**Fig. 4.**
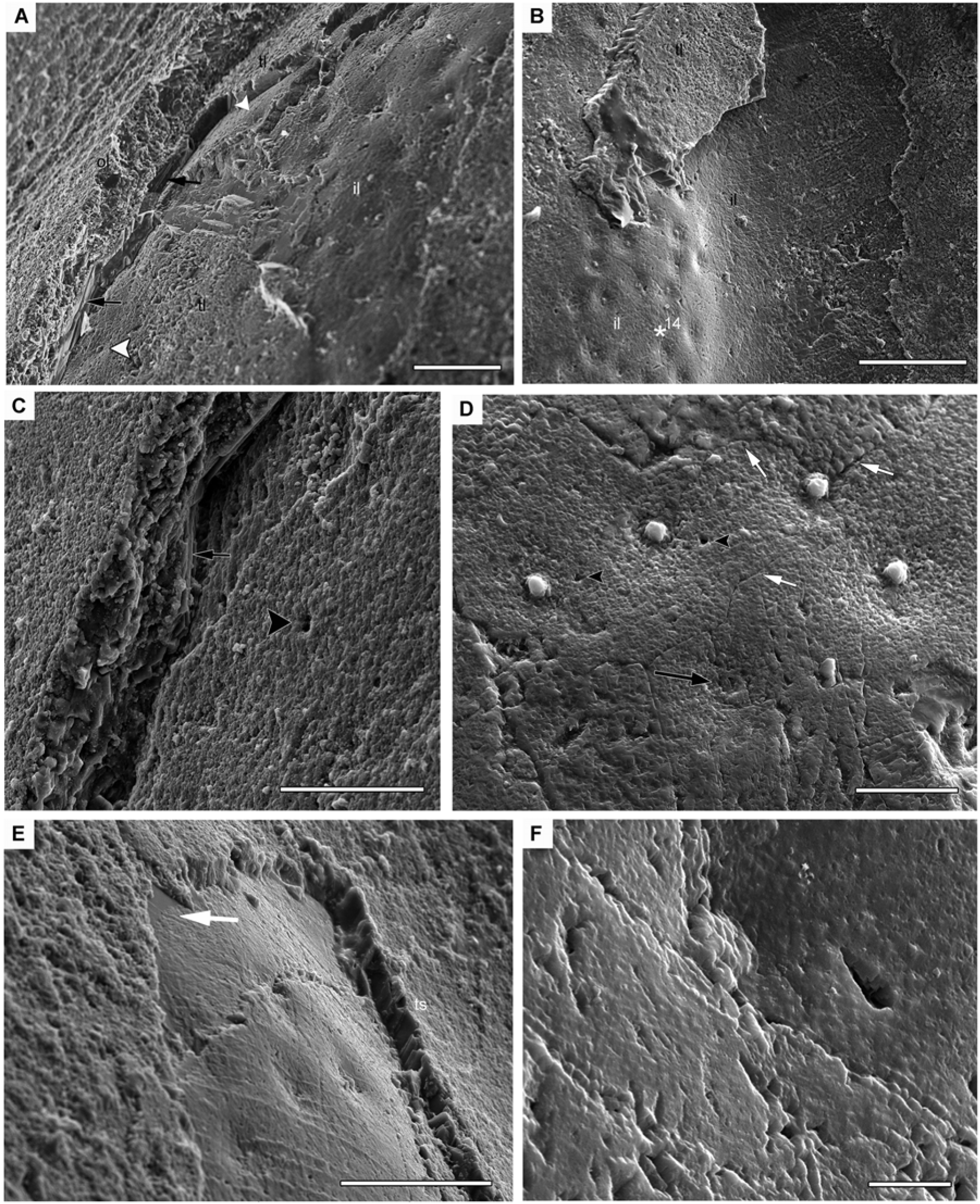
Cephalon *Pseudagnostus sp. ex gr. P. cyclopyge* PIN 5862/1. A, a fragment with the clearly visible layered architecture of the cuticle: relatively loose upper layer with the outermost polygonal pattern, dense transitional layer showing horizontal (here) lamination (black arrows), dense inner layer with peg pits (white arrowhead), and basal subunit of the inner layer with peg pits. B, the fragment with a piece of the transitional layer (vertical lamellae are visible as columnar structures at its left fracture plane), the inner layer with peg pits and the rhombic striation in the furrow. C, a fracture showing the comparatively loose outer layer where the horizontal lamination can be supposed, the transitional layer displaying thin horizontal lamination (black arrow), the double pore is clearly discernible on this surface (black arrowhead); the rhombic mesh is visible under the laminated transitional layer, the surface of the inner layer with peg pits is not displayed in this fragment. D, the surface with the peg pits: pegs with the whitish tops are visible, one pit lacks the cone, the traces of a slightly convex collar with an indentation inside substitutes the peg (black arrow); pores are present on the surface (arrowheads); very thin fractures (white arrows), which repeat the outermost polygonal pattern, are visible too. E, a fragment of the transitional layer with vertical lamination (see also Fig. 3M) enclosed a patch of the inner layer where the rhombic mesh and peg pits are visible; the left corner of this patch shows its thinning to the host rock (white arrow) demonstrating the width of the inner layer (less than 7 μm). F, the enlarged fragment of the basal subunit with a pit and the base of a peg; tiny regular indentations as well as rhombic mesh cover the surface, both are possibly reflecting a network of chitin fibrils of the subunit. Scale bars: 50 μm (A, C, E); 100 μm (B); 20 μm (D); 5 μm (F).

**Fig. 5.**
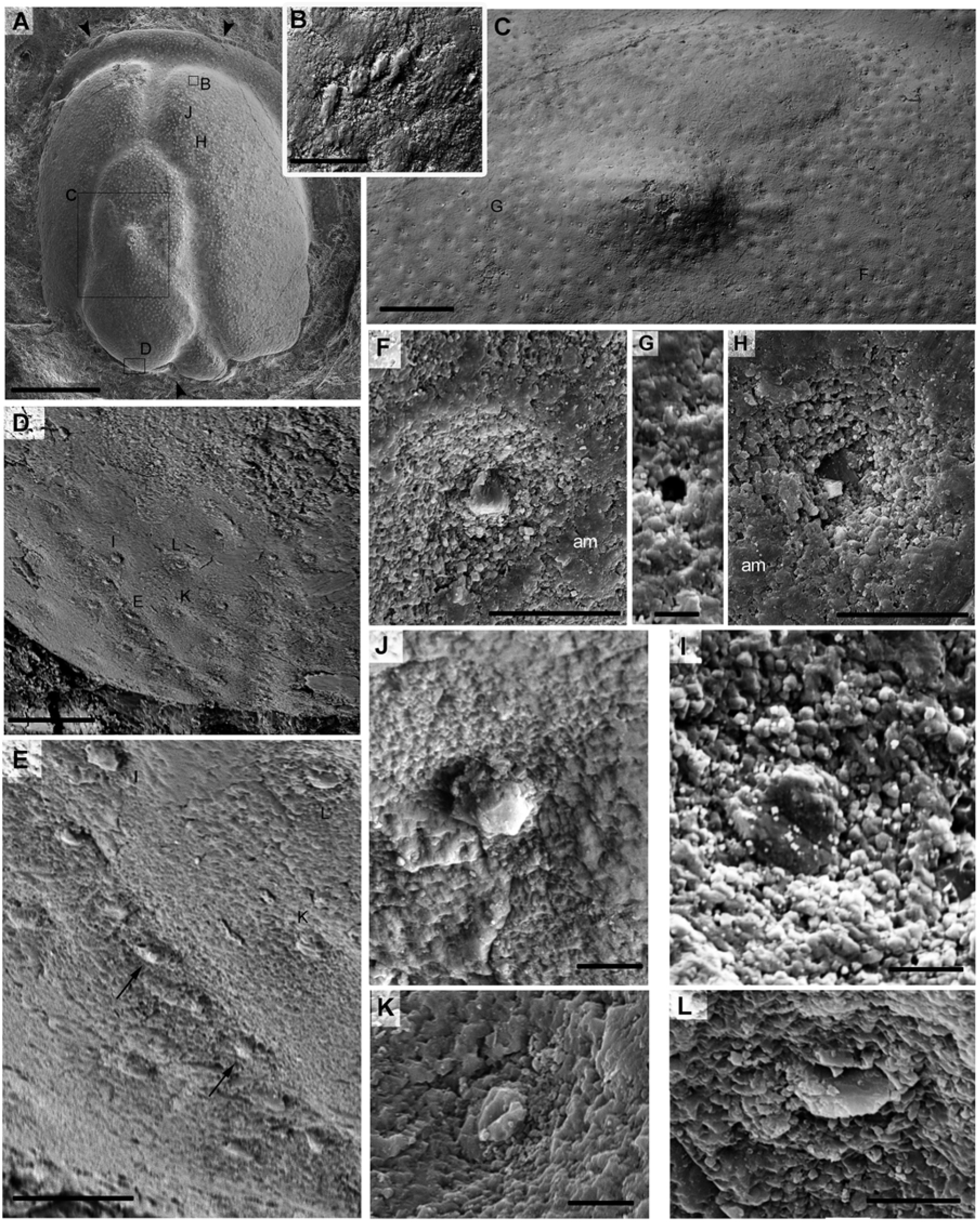
*Pseudagnostus sp. ex gr. P. cyclopyge* PIN 5862/2. A, general aspect; arrowheads show fragments of the outer layer, the rest of the surface represents the inner layer covered by numerous peg pits visible as dots. B, fragment with four parallel flat elongate tubercles on the inner layer of the right acrolobe (the symmetrical fragment is destroyed). These tubercles resemble placoid sensilla. C, fragment of the glabella with a glabellar node and muscle scars displaying the inner layer with evenly distributed peg pits; they are few on the muscle scars. D, the glabellar culmination with a V-shaped groove; rows of narrow elongate tubercles located parallel to the glabellar base are clearly visible; these tubercles resemble the shape of digitiform sensilla. Several other relatively large elongate tubercles (shaped as campaniform sensilla) are situated in the vicinity of the groove and between its branches. E, the left branch of the V-shaped groove where the elongate tubercles within the groove branches are exposed; in some of them small convex bosses on top are visible (black arrows). F, typical shape of a peg pit: a shallow depression with a central roundish peg. Some amorphous material (non-crystalline or very fine crystalline; *am*) can be seen around the pit. G, an example of rimless pores between peg pits. H, a regular pit without a central peg; there is an opening instead of it (the nature of the whitish crystal in the opening is unknown). I, a peg in the pit lacking any whitish top cover, there is a tiny opening on the top of the peg. J-L, there are several types of tubercles surrounding the V-shape groove: J, a pit with a low pentagonal cone which is surrounded by a collar; it can represent a stiloconic sensillum. K, a collar with an indentation; the peg of some other element was possibly destroyed. L, an elongate tubercle surrounded by a collar; this is the only tubercle of all others around the V-shaped groove displayed both on the outer and inner layers. This is possibly a campaniform sensillum. Scale bars: 1 mm (A); 50 μm (B, D); 200 μm (C); 20 μm (E, F, H); 4 μm (G); 5 μm (I–L).

In both specimens, the outer layer is 30-40 μm thick. Its outermost subdivision with polygons is the thinnest and easily exfoliates (Figs 3B, I). In some areas, its crystal structure is clearly visible: it is composed of densely packed small prismatic crystals (Fig. 3G). These crystals are unlikely to be the result of post-mortem recrystallization or post-mortem deposition on original cuticlar biominerals, as we can see rims of regular crystals around the pores (Fig. 3H) which are definitely original biominerals.

The principal part of the outer layer appears to be composed of horizontal laminae which can be evenly split one from another (Fig. 3B). On the fracture planes, it can sometimes be seen that the principal part is composed of relatively thick, loosely packed horizontal lamellae (Fig. 3D, arrow).

Pore openings protrude onto the surface of the outer layer and are reinforced with crystalline rims (Figs 3G-J). We also managed to see the canal traversing horizontally through the outer layer (Fig. 3E). Its wall is comparatively smooth, with minute, ordered pits (Fig. 3F), resembling in appearance the surface of the basal sublayer (Fig. 4F).

In the glabellar culmination, there is a V-shaped groove with a row of bulbous tubercles (Figs 3C). A convex oval element is located in the center between the branches of the groove (Fig. 3C (arrowhead), Fig. 12M).

The underlying transitional layer has a simple surface, sometimes with vague traces of the peg pits which are specific elements of the underlaying inner layer (Fig. 3N). It also bears multiple pores but without a crystalline rim (Figs 3K, 4C).

This layer is built in stacks of horizontally, vertically or obliquely laid thin lamellae (Figs 4B, D, L, M, O). One of the fracture planes shows the connection of two stacks with lamellae directed differently (Fig. 3O). In the other parts, the lamellas are directed horizontally (Fig. 4C) or vertically or obliquely (Fig. 3M). The lamellae of this layer are thinner than the lamellae of the outer layer.

The transitional layer superimposes the inner layer. On the cephalon 5862/1, the layers split exactly along the boundary of the inner layer and the upper layers. The surface of the inner layer differs sharply in appearance from these two: there are numerous regularly shaped peg pits on the inner surface. The maximum inner layer thickness is less than 7 μm (Figs 3L, 4E).

In the specimen 5862/2, remnants of a non-crystalline (or very fine-crystalline) substance are visible on the surface of the inner layer between the pits (Figs 5F, H, 6D, G, F). On the border, there are convex structures embedded in this substance (Fig. 6H). This substance delimits the inner layer from the overlying cuticular subdivisions.

**Fig. 6.**
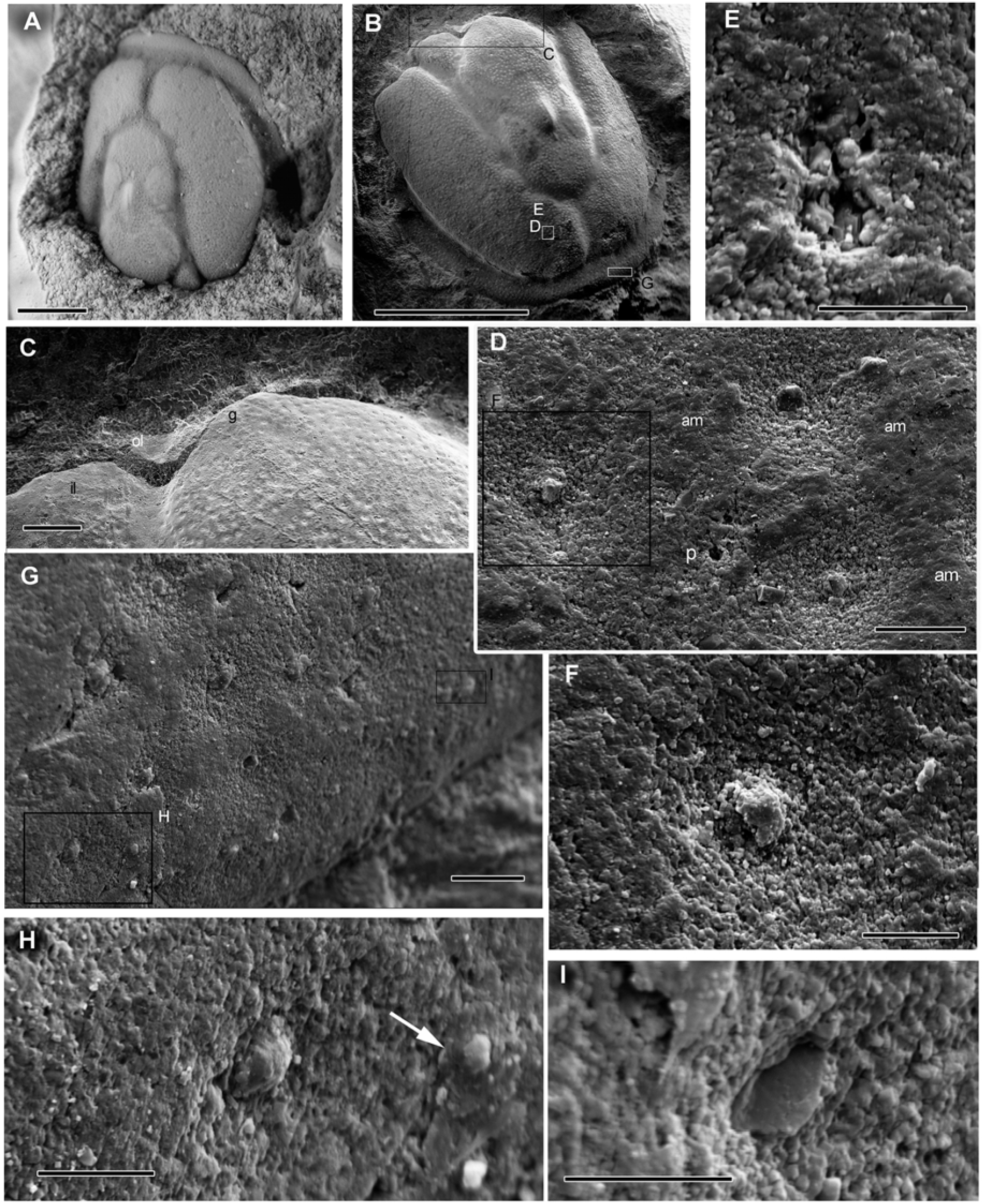
*Pseudagnostus sp. ex gr. P. cyclopyge* PIN 5862/2. A, general aspect of the specimen under an optical microscope with magnesium coating. The specimen was imaged before the preparation of the cephalic rear and the left side of the acrolobe. B, general aspect of the specimen under SEM showing regularly scattered peg pits. C, glabellar culmination showing the fragments of the outer layer and the V-shape groove (g). D, a typical pattern of the inner layer with peg pits: the surface is covered by an amorphous material (am) except for the pits, the pegs have some whitish tops, round pores (p) are situated between the pits. E, a double pore between pits; the crystals inside the pores can be either of biological or depositional origin. F, typical pit with a peg covered by whitish material. G, a fragment of the border showing multiple peg pits and convex tubercles. H, two oval convex tubercles, the right one is imbedded into some amorphous material (white arrow) with its top clear of it. I, an example of the elongate tubercles in the border showing a thin rim around the tubercle. Scale bars: 1 mm (A); 500 μm (B); 200 μm (C); 20 μm (D, G); 10 μm (E, H, I); 5 μm (F).

Very thin fractures corresponding to the polygonal pattern can be seen on the surface of the inner layer (Fig. 4D, white arrows). Also, the traces of rhombic striation from the underlying basal subunit are sometimes exposed (Figs 4B, F).

The basal subunit of the inner layer has a surface with a rhombic mesh (Figs 3D, M, O, P; Fig. 4C). Its thickness is about 0.5-1 μm. It is strictly ordered and can be seen both explicitly (Figs 3M, O, 4C, E) and as a rhombic striation (Figs 3D, P, 4A, B). On the surface of this sublayer, the peg pits are visible, but the pits are quite shallow, and the cones are expressed only as low bosses (Figs 3O, 4E, F).

On the inner layer the V-shaped groove was also distinct. Here, the groove looks like a continuous depression instead of a series of pits on the outer layer. In the symmetrical branches of the groove, there are elongated tubercles located parallel to each other and to the base of the glabella. In the diverging upper part of the branches, the tubercles seem rounded or oval, while in the lower part they are elongated. A relatively large convex oval tubercle in the shallow depression is located between the branches on the symmetry axis (Figs 5E, L).

The two studied cephalons differ in their chemical composition (Table 1). In the cephalon 5862/1, Ca prevails in the outer layer with a small supplement of Mg while in the inner layer Ca is significantly lower. There is about the same amount of Ca in the inner layer as in the surrounding rock. The supplement of Mg in the inner layer is recorded only in the surface amorphous/fine crystalline substance. However, there is a noticeable amount of Al and Si in the host rock, but these elements are practically absent both in the outer and in the inner layer.

In the carapace 5862/2, Ca is abundant in the inner layer, whereas Al and Si are there only in negligible amounts. However, the outer layer does not differ in composition from the host rock in any of the studied elements.

From the contrast in the elemental composition of the cuticular layers and the surrounding rock, it is easy to understand that both these layers are not casts or imprints of the cuticle on the host rock. Meanwhile, the outer layer of cephalon 5862/2 has the same elemental composition as the host rock. However, this layer cannot be some kind of postmortem deposit as there are definite morphological features (that is, the polygonal pattern, the pores with the rims, the V-shaped groove, loose horizontal lamellaes). Thus, we can assume that a complete replacement of this cuticular layer took place.

### Pseudagnostus josepha (Hall, 1863) (Fig. 7)

This species is represented by a cephalon and pygidium from the same bed.

The cephalon possess small pieces of the outer layer (Figs 7A, G). As can be seen on one of the fracture planes, this layer is composed of a stack of horizontal lamellae (Fig. 7J). Small openings or indentations (1μm) are visible on the outer layer (Figs 7 L, M). Sometime rather complex structures could be seen on the outer layer like those in Fig. 7K.

**Fig. 7.**
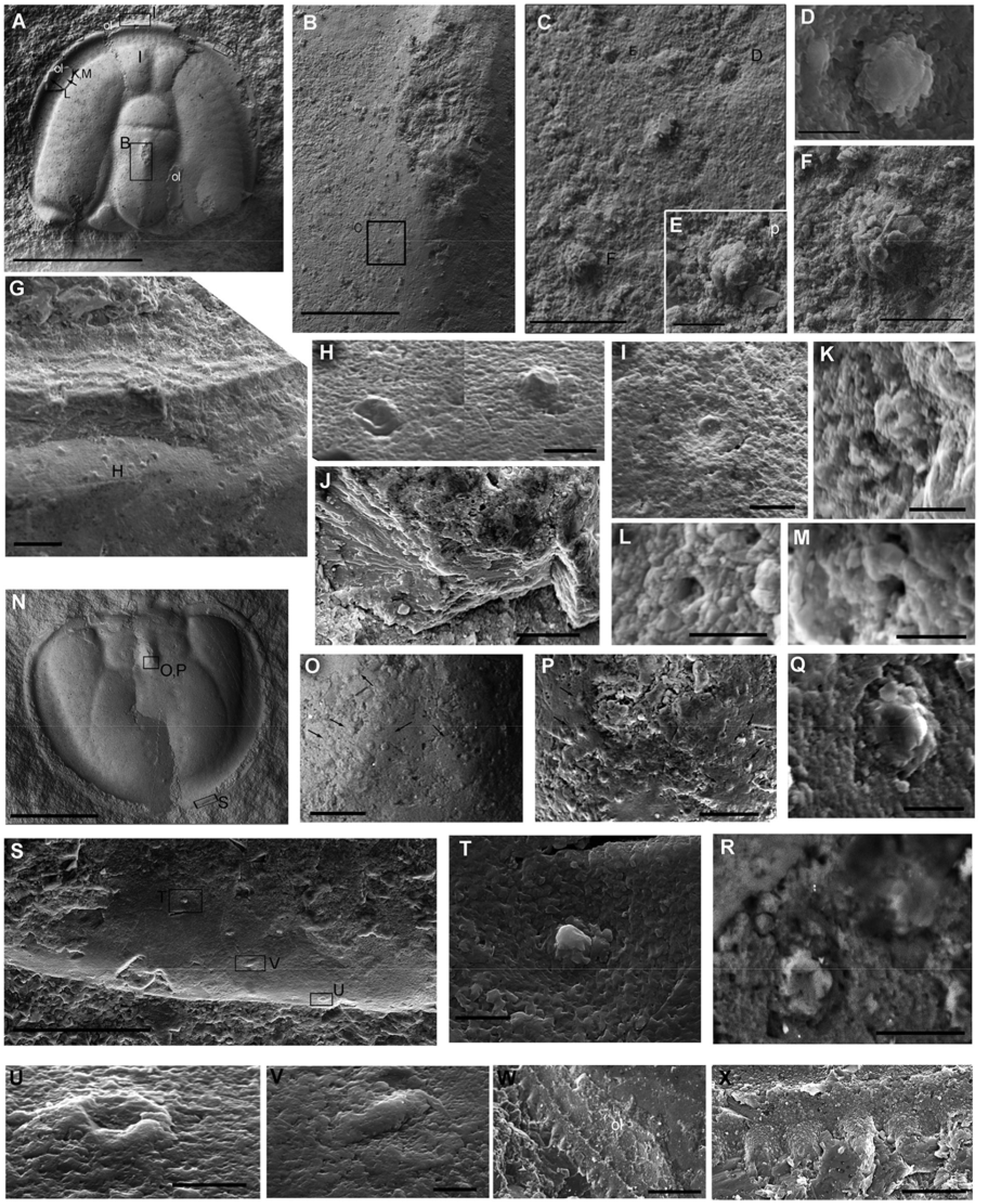
*Pseudagnostus josepha* (Hall, 1863) A–M, cephalon PIN 5862/11. A, general aspect of the cephalon; small pieces of the outer layer (ol) are visible. B, the surface under the outer layer with scattered tubercles (or their casts on the rock); we suppose that this is a replacement of the inner layer or a transitional one. C, boss-like tubercles (or their casts) and a shallow pit with a central structure (enlarged in D). D, a complex element in the center of the pit; it is larger than the typical pegs in the pits. E, a boss-like tubercle and a very small pore (p) next to it. F, the comparatively large boss is built from the clay crystals that allow us to suggest a diagenetic transformation of this layer and/or the cast nature of it. G, fragment of the outer layer on the border with preserved polygonal ridges on the surface; there are multiple tubercles of different sizes and shapes underneath the outer layer. H, two examples of convex tubercles with smooth surfaces: the left one is elongate and probably crinkled, which indicates the original elasticity of the structure, the right one has a round shape with a shallow depression in the center. I, pit with a central cone; most of these pits are smoothed on this surface. J, a fragment of the outer layer with a visible thin lamination. K–M, structures on the small piece of the outer layer: K, a flower-like element with a tiny central opening. L, an indentation with a central base in its bottom. M, a small rimmed pore. N–V, pygidium 5862/12: N, general aspect of the pygidium; the outer layer was not preserved in this specimen. O, an area behind the axial node showing at least four rows of some convex round elements (black arrows) situated symmetrically on either side of the axial node. P, the area with these rows; the convex shape of these elements are well visible. Q, an example of these elements demonstrates the complex shape with the flower-like base. R, two other examples of these elements showing the flower-like base. S, a fragment of the border preserving multiple tubercles and peg pits. T, a typical peg pit. U, convex elongate tubercle with an indentation. V, convex elongate tubercle. W-X, pygidium *P. ampullatus PIN* 5862/13: W, edge of the border between anterolateral spines, which bears a row of tubules (or blunt spines) on the outer layer. X, enlarged tubules (spines). Scale bars: 2 mm (A, N); 300 μm (B); 100 μm (S); 50 μm (O, P, W); 25 μm (C); 20 μm (G, X); 10 μm (F, I, R); 5 μm (D, E, H, K – M, Q, T, U, V); 3 μm (J).

The illustrated pygidium (Fig. 7N) did not preserve any fragments of the outer layer.

The inner surface of the cephalon and pygidium looks rough (Figs 7B, G, S). The roughness is composed of tubercles and bosses of various sizes and shallow depressions with simple or stellate tubercles (Figs 7C-F). The borders look smoother (Fig. 7 G, S) however they possess a lot of various protuberances (Figs 7H, U, V). Openings are visible in this surface. Well preserved peg pits also present on the cephalic and pygidial borders (Fig. 7T), and poorly preserved ones were found on the cephalic acrolobes (Fig. 7I).

The prominent part of the pygidial axial node is surrounded by interesting bulbous bodies (Figs 7O, P). They are arranged in three-four symmetrical rows that go down from the node. In some of these bodies, a star-like base with a convex top are distinguished (Fig. 7 Q, R). These bodies are of around 7 μm.

EDX-point analyses of the cephalon (Table 1) revealed differences in the elemental composition of the outer and inner layers. The outer layer contains a high amount of Ca with a minimal addition of Al and Si compared with the surrounding rock. The average elemental content of the inner layer is more or less the same as the rock; however, the dispersion is quite large. Some patches and microstructures in the inner layer are mostly calcareous (points 7-9 and some other ones) while others resemble the elemental composition of the rock.

### Machairagnostus sp. (Fig. 8)

This small cephalon (Fig. 8A) displays two cuticular subdivisions (Fig.8 B). The width of the outer layer is 10 – 20 μm (Figs 8B, G). The outermost surface possesses a distinct ridged polygonal pattern (Fig. 8C). The polygons are preserved very well in the front part of the acrolobe but gradually disappear around the glabella. This shows that the polygons can be easily destroyed and smoothed in a fossilized specimen. The principal subdivision possibly consists of loose horizontal laminae which are sometime seen in fracture planes (Figs 8B, G).

**Fig. 8.**
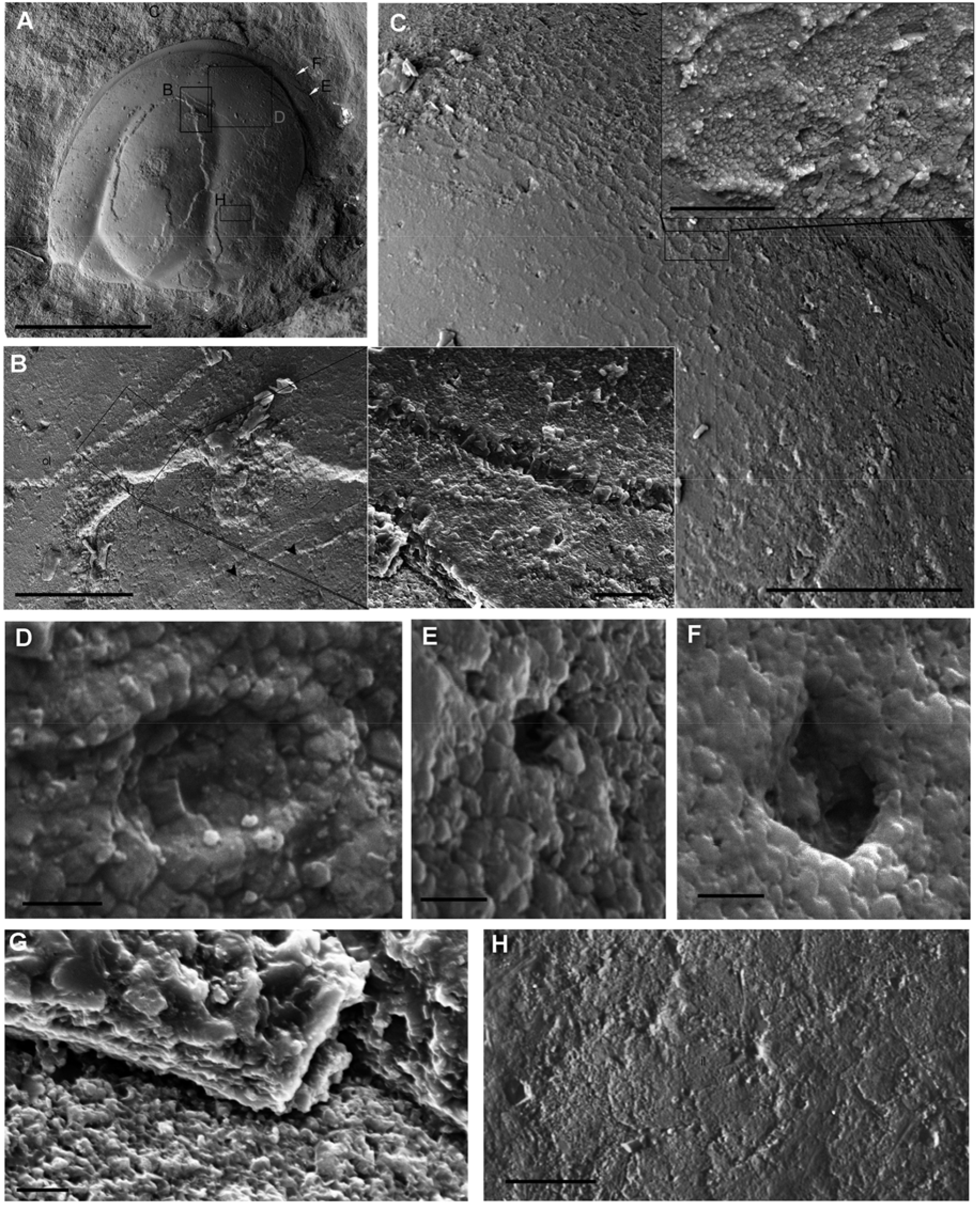
Cephalon *Machairagnostus sp*. PIN 5862/4. A, general aspect; an outer layer (with two sublayers) is visible on the acrolobe, basal lobes, and border while the central part of the glabella lacks it. B, two sublayers of the outer layer (see inset on the right: outermost prismatic sublayer and loosely laminated principal sublayer are well visible); black arrowheads show thin polygonal fractures fractures on the surface under the outer layer. C, a fragment of the outer with the polygons which smoothed out to the glanellar front; in the inset examples of the ridged polygons. D–F, structures on the surface of the outer layer: D, a framed indentation with the central opening surrounded by very thin walls. We supposed that this wall is the remain of some longer hollow seta. E, deep indentation (possibly a pore). F, indentation with a central tubercle on its bottom. G, fracture plane of the outer layer showing its loose texture with possible horizontal lamination. H, surface under the outer layer with very thin fractures arranged as polygons; it could be either traces of the epithelial layer or an inner cuticular layer. Scale bars: 1 mm (A), 100 μm (B, inset 20 μm); 200 μm (C, inset 20 μm); 2 μm (D–F); 5 μm (G); 50 μm (H).

There are comparatively small pores (1-1,5 μm) on the outer surface. Also, different indentations of various shapes were found on the surface of the acrolobe and border (see below) (Figs 8D, F).

Under this layer, there is a surface bearing very thin fractures arranged in the polygonal pattern (Figs 8B, H).

All layers display more or less the same elemental composition as in the host rock matrix (Table 1). However, the outer layer contains increased Ca and decreased Mg, and much less Si and Al. The inner layer shows decreased Ca, Mg, and Si compared to the rock. This difference allows us to suppose that this surface represents the inner cuticular layer or specifically, a replaced hypodermal cell layer rather than a cast of the cuticle on the rock surface.

### Aspidagnostus lunulosus (Kryskov, 1963) (Fig. 9)

The cephalon and pygidium of A. lunulosus came from different beds (Figs.1). Both have patches of the outer layer (Fig. 9A, F, B, G), however we did not detect any surface polygonal pattern on it, nor pores or other organized structures.

**Fig. 9.**
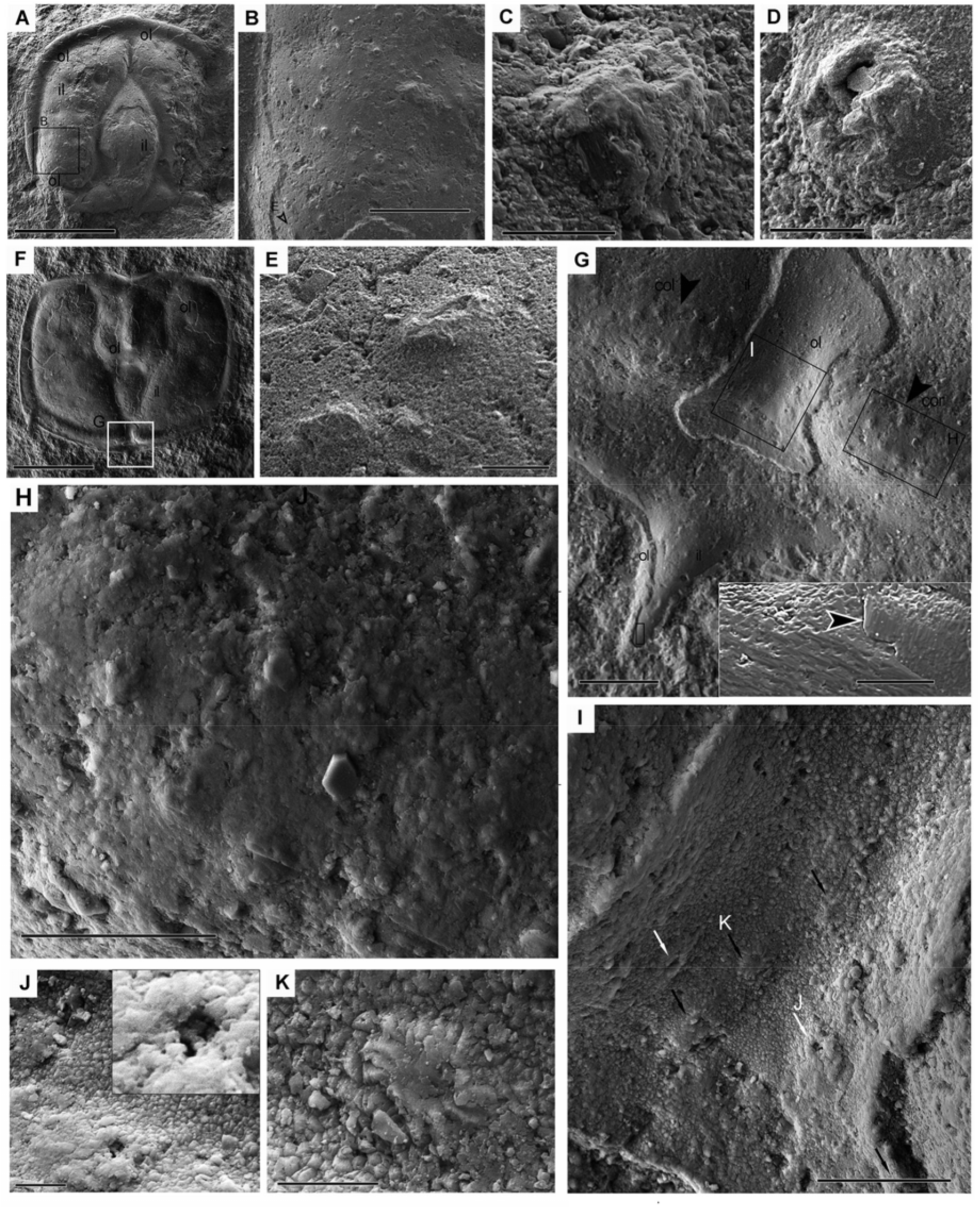
*Aspidagnostus lunulosus* Kryskov, 1963. A, general aspect of the cephalon PIN5862/14. Patches of the outer layer (ol) are visible with the inner layer (il) below. B, enlarged fragment of the inner layer (the outer layer is visible on the sides) showing numerous elongate and round tubercles. C, D, round tubercles with the central indentation. E, two elongate tubercles. F, general aspect of the pygidium PIN5862/15. Patches of the outer and inner layers are visible. G, zonate border with the piece of the outer layer in the postaxial furrow, fragments of the outer layer are also preserved around the central spine. The elliptical rows of elongate tubercles in the right (cor) and left (col) parts of the border collar are clearly visible (black arrowheads). A group of indentations in the postaxial area can be also discerned. Inset: the surface of the inner layer in the central spine; there are numerous tiny indentations spread regulary upon the surface, the joining of two plates of unequally directed laminae or fibrils (black arrowhead with white outline) are shown. H, enlarged fragment with the elliptical row of elongate tubercles in the right part of the zonate collar. I, enlarged fragment with a group of indentations in the postaxial area (shown by arrows). J, K, the best-preserved examples of the right-hand (J) and left-hand (K) structures of I. Scale bars: 1 mm (A, F); 200 μm (B); 100 μm (G, inset 10 μm); 50 μm (H, I); 20 μm (E); 10 μm (C, J, K); 5 μm (D).

On the inner cuticular layer of the cephalon, there are numerous tubercles of different shapes and sizes (Fig. 9B). We found elongate tubercles (Fig. 9E), relatively large round tubercles with indentations (Fig. 9C, D) and small ones without indentations (not shown here).

The inner layer in the pygidium does not carry pronounced tubercles like those on the cephalon. There are few small bosses on the border and central border spine (visible in Fig. 9G at magnification). The inner layer is possibly composed of a network of thin laminae obliquely superimposed onto each other; the surface of these laminae displays regular tiny indentations (Fig. 9G, inset) the same as in the basal fibrillary sublayer in *Pseudagnostus sp. ex gr. P. cyclopyge* (N5862/1, Fig. 4F). There is a spectacular suit of microstructures on the rear part of the zonate border and postaxial furrow. Firstly, in the postaxial furrow, a piece of the outer cuticular layer was preserved (Figs 9F, G). It has at least three seemingly identical elements located in the bottom of the postaxial furrow (Fig. 9I, black arrows). The best preserved one looks like a shallow deepening in the center of a rim of elongate prismatic crystals (Fig. 9K). In the axial furrow on either side of these prismatic rimmed structures, there are at least two symmetrical openings or indentations with some central elements inside (Figs 9I, J).

On the right and left collars of the zonate border, there are symmetrical groups of identical elongate tubercles arranged along ellipsoidal curves (Figs 9G, H). No morphological details of these tubercles were clear due to poor preservation. This suit of regularly organized structures both on the collar and the axial furrow is most probably a representation of some sensorial ensemble.

Elemental analysis of the pygidium indicates an increased amount of Ca in both the inner and outer layers in comparison with the rock around the specimen (Table 1). Mg appeared to be slightly lower in the outer layer than in the inner layer and in the rock. Al and Si are present in all samples, Si being higher in the rock. Therefore, the elemental compositions of the specimen confirm that this is the cuticle rather than a cast in the rock.

### Homagnostus sp. (Fig. 10)

This cephalon bears remains of the outer layer on the left side of the acrolobe (Fig. 10A). Its width is about 6-10 μm and it bears a distinct reticulate pattern shaped out by ridges. The outermost surface is prismatic, and the middle and bottom part of the layer is probably composed from several (at least 15) thin lamina (Figs 10B, C). Elongated convex bodies are located between ridges (Figs 10C). Also, there are rimmed pores on this surface (Figs 10D, E).

**Fig. 10.**
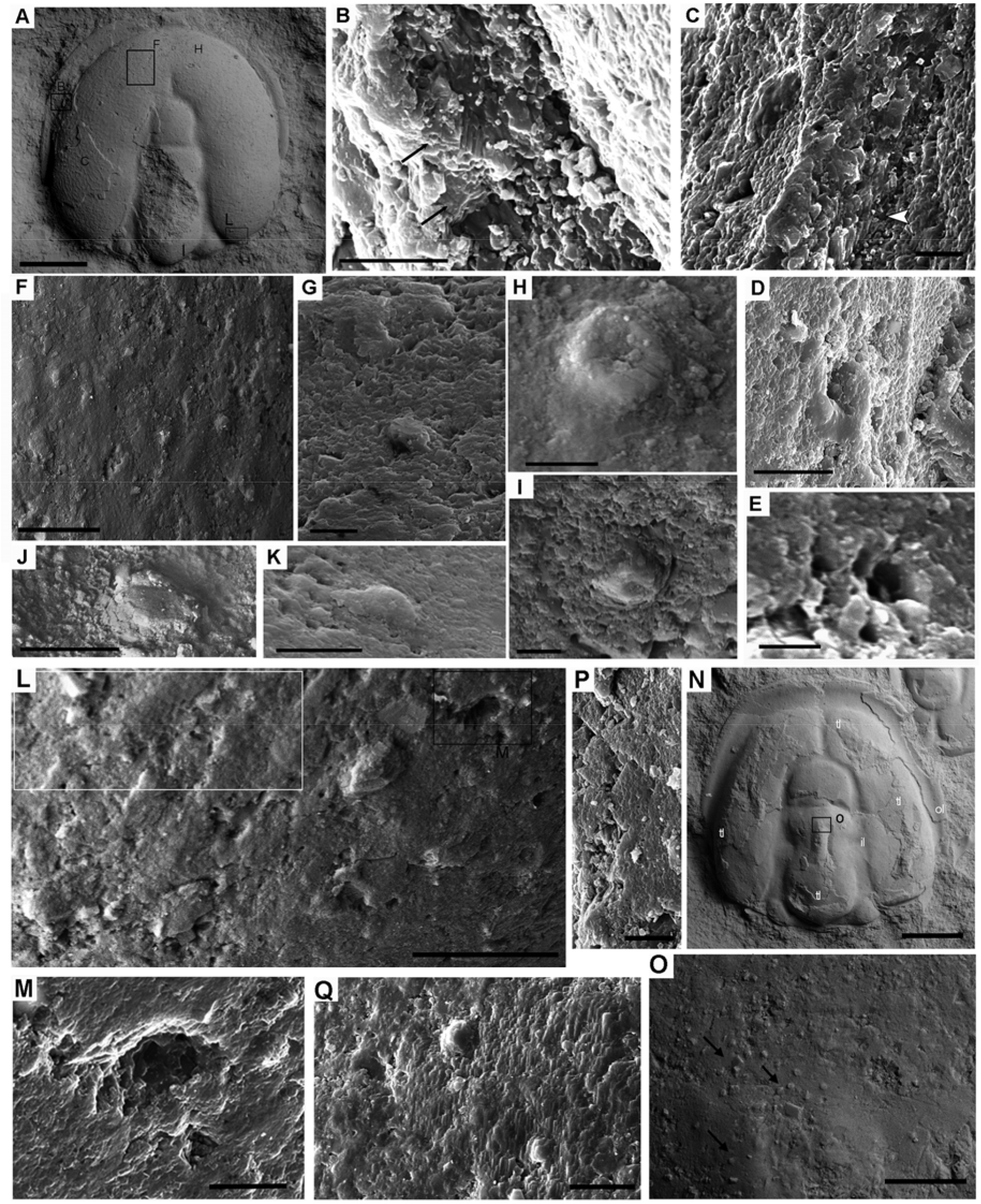
Cuticular structures in Agnostidae and Ammagnostidae: A–M, *Homagnostus sp*. PIN5862/15: A, general aspect; fragments of the outer layer with the polygonal pattern are visible on the acrolobe at the left. B, enlarged fragment of the outer layer with a thin lamination discernible in the angular fracture (black arrows). C, crystalline surface of the outer layer with polygonal ridges and a convex smoothed elongate element between the ridges; there are thin horizontal lamellae visible at the lower part of the outer layer (white arrowhead). D, a rimmed pore in the polygonal surface; the rim is covered by non-crystalline or very fine crystalline material. E, a small rimmed pore; the rim is built of comparatively large crystals; the polygonal pattern is lacking on this fragment of the outer layer. F, surface of the inner layer; traces of the some rhomboid relief can be recognized in it; convex elements are present. G, shallow pit with a central element on the inner surface. H, I, examples of convex bosses with a central depression. J, K, elongate (rhomboidal) convex bodies which represent the most numerous elements on the inner layer. L, the surface with the convex elongate bodies and a pore; the framed rectangle shows a patch where a meshed structure of the inner layer is discernible. M, enlarged fragment with the funnel pore in the inner layer; it possibly has a meshed bottom. N–P, cephalon *Proagnostus bulbus* PIN5862/16: N, general aspect, fragments of the outer layer with the polygons are clearly visible; meandering pattern on the transitional layer is visible under magnification. O, small bosses around the top of the median glabellar node (black arrows) arranged possibly in rows. P, rhomboid plates composed the inner layer. Q, two pits with the central structures. Scale bars: 500 μm (A); 20 μm (B); 5 μm (C, E, G – I) 10 μm (D, J, K, M, P, Q); 50 μm (F, L, O); 1 mm (N).

The inner layer shows traces of the surface reticulation seen as a low rhomboid relief; small round casts or bodies located within these rhombuses (Fig. 10F). Round convex bosses with shallow indentations in the center are also present in the inner surface of the front and middle parts of the acrolobe (Figs 10H, I). Larger rhomboidal bodies are located regularly in the rear part of the acrolobe (Fig. 10L). Conversely, we found a few pits with the central structure on this surface however all of them were smoothed in this specimen (Fig. 10G). Along with the various convex elements there are comparatively large openings (8-10 μm) without rims which are easily distinguished from any surface breakages (Fig. 10 M).

Elemental compositions of the outer and inner layers show considerably (2-2,5 times) increased Ca compared to the rock while Mg was 3-3,5 times lower in the outer layer than in the inner layer and in the rock. Si and Al complement the composition of the rock and the shell, though Al in the inner layer was considerably (2,5 times) decreased (Table 1).Since the elemental compositions of the inner layer and the rock were different, the inner surface can be supposed not to be a cast on the rock.

### Proagnostus bulbus (Figs 10N-P)

The imaged cephalon (Fig.10N) is representative of a number of specimens *of P. bulbus* on a single slab. They all have the same morphology of the cuticular structures. Their cuticle consists of at least two layers. The fragments of the outer layer (25 μm in width) on the imaged cephalon are easily distinguished by reticulate surficial pattern. This reticulation seems to be smaller and more regular in the border than in the acrolobe. A vertical chip across the outer layer clearly shows that the outermost sublayer is extremely thin (*c.*1μm) and differs in texture from the rest of the outer layer (not shown here). The outer layer is loosely adjacent to the inner layer: a very thin gap is visible between them. No distinct elements (pores or tubercles) were found on the outer layer.

A transitional layer is possibly present here: it manifested itself in the meandering ridged pattern or its counter-cast on the surface (Fig.10O).

The inner layer is very thin (less than 3 μm). Shallow pits with a central boss are sometime discernible on this layer (Fig. 10N).

In the inner layer, a group of interesting structures was found to surround the median glabellar node. This group is represented by small round bosses (3-4 μm) arranged in two or more rows going down from the top of the glabellar node (Fig. 10Q). It should be mentioned that we noticed the same group in several other specimens, and in one specimen, instead of bosses, there were a group of indentations with some central structures inside.

The elemental compositions of the rock, outer, transitional, and inner layers appeared to be more or less similar (Table 1) and thus we could not distinguish whether it was a cast or true cuticular layers. Because of this, the chemical composition did not help us to infer if the meandering pattern on the transitional layer and the interesting group of bosses around the glabellar node represent the remains of some anatomical structures, or if it is a cast of the visceral cuticular surfaces.

On the inner layer the V-shaped groove is displayed. Here, the groove looks like a continuous depression instead of a series of pits on the outer layer. In the symmetrical branches of the groove, there are elongated tubercles located parallel to each other and to the base of the glabella. In the diverging upper part of the branches, the tubercles seem rounded or oval, while in the lower part they are elongated. A relatively large convex oval tubercle in the shallow depression is located between the branches on the symmetry axis (Figs 5E, L).

In this section, we discuss the following issues: (1) the divisions of the agnostoid cuticle; (2) comparison of the cuticular arrangement in agnostids and in classes of arthropods; (3) cuticular pores and sensilla in agnostids.

## DISCUSSION

### The arrangement of the agnostoid cuticle (Fig.11)

In the cuticle of the studied specimens, three layers can be discerned (Fig. 11). Depending on preservation, only part of the full set of the cuticular layers and their subdivisions are displayed. The cuticular architecture traced most completely in the carapace of *Pseudagnostus sp. ex gr. P. cyclopyge* PIN 5862/1.

**Fig. 11.**
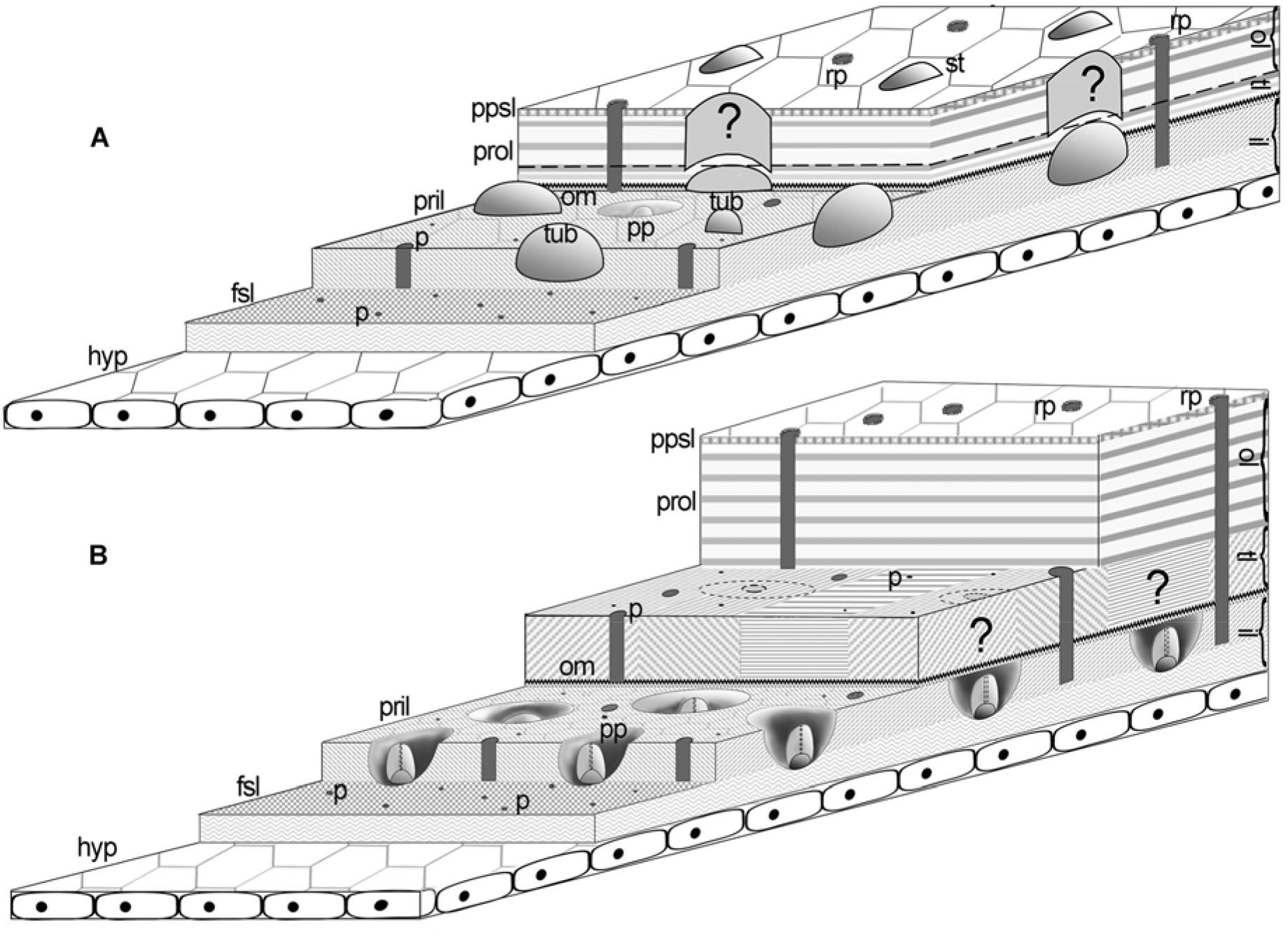
Architecture of the agnostoid cuticle. A, Agnostidae, Ammagnostidae, Aspidagnostidae; B, Pseudagnostidae; lettering for layers: hyp – hypoderma, ol – outer layer, tl – transitional layer, il – inner layer, fsl – basal fibrous sublayer, om – rich organic interlayer space, pril – principal inner layer, prol – principal outer layer, ppsl – polygonal pattern sublayer; lettering for microstructures:, pp – peg pit, p – pores, rp – rimmed pores, tub – convex tubercles of different shapes; shadings: horizontal, oblique and vertical – horizontal, oblique and vertical lamination, cross hatching – fibrous chitin-proteinaceous mesh.

#### Outer layer

The outer cuticle layer is subdivided into two units. The uppermost unit bears the surface polygonal pattern; it is very thin and possibly built of prismatic crystals. It is easily erased and thus appears as ridged patches. This kind of reticulation is well known – for example, it was studied in crustaceans (Giraud-Guille 1984) and horseshoe crabs (Mutvei 1981). It has been shown that while biomineralization of the carapace proceeded, intercellular septa became mineralized first and more heavily, and due to this, polygonal ribs were formed (Dillaman *et al*. 2005). Presumably, agnostids did not differ from other arthropods in this feature, and the polygonal pattern reflects the shape and size of the hypodermal cells producing biominerals. The shape of the polygons seems to change in the direction of increased growth or elongation of hypodermal cells that secrete the outer layer (Miller 1976; Müller and Wallossek 1987; Wilmot 1990, Dalingwater *et al*., 1991; Giraud-Guille 1984). In our material, the smallest and most elongated polygons were observed at the dublura in pseudagnostids. It is likely that skeletal-producing cells grew most actively there.

The principal part of the outer layer has a loose lamellar structure. Where the lamellar structure was visible, the laminas were horizontal. Since lamellae in the outer layer were found in representatives of two families (Agnostidae and Pseudagnostidae), the loose-lamellar structure of this layer is most likely a common feature of agnostids. It should be noted that the lamellar structure in the outer layer is difficult to detect as we found it in only a few fracture planes, and previous studies failed to find any signs of lamination.

The outer layer varies greatly in thickness, ranging from 10-30 μm in adult specimens of Agnostidae and Ammagnostidae and up to 30-100 μm in adult specimens of Pseudagnostidae. It is not however known whether the thickness of the outer layer varies on cephalons and pygidia, as well as in their different areas.

#### Transitional layer

This layer was clearly identified in the specimens of *Pseudagnostus sp. ex gr. P. cyclopyge* (PIN5862/1) and in two pygidia of this species described elsewhere (Naimark, Chaika, 2022) and *P.bulbus* PIN5862/16. The thickness of the *Pseudagnostus sp. 1 ex gr. P. cyclopyge* was 12-20 μm, and of the *P. bulbus* specimens was 8-15 μm (one and a half times less than in the outer layer in this specimen).

Like the outer layer, it might be composed of lamellae, but here the lamellae were thinner than those in the outer layer. The lamellas appeared to be packed in stacks; in each stack the lamellae sloped differently. In *P. bulbus* the transitional layer bore some meandering surface pattern, while in *Pseudagnostus sp. ex gr. P. cyclopyge* there was not any surface pattern. Instead, there are traces of the peg pits which were located in the underlying inner layer.

In the two studied pygidia and several other cephala of Pseudagnostidae, the transitional layer does not split out or distinguished in appearance from the outer layer in the fracture planes. Only in single fragments could we recognize its presence by the specific traces of the peg pits on the patches of the surface splitting from the outer layer. Thus, we could conclude that this layer tightly adjоins to the outer one and can possibly merge into it in some areas.

In many parts of the cuticle, a very thin gap is clearly detectable between the outer/transitional layers and the inner layer. Thus, the splitting along the visceral surface of the outer/transitional layers and the inner one seems quite natural. The origin of this gap is not yet clear; presumably, it could be filled with organic matter or a biomineralization process changed at this particular boundary. In any case, the phenomenon of splitting along this surface indicates the specificity and integrity of the two upper layers and the inner layer.

#### The inner layer and its basal fibrous sub-layer

The inner layer is quite thin (>7μm). It has a specific surface distinct from the two upper layers: in Pseudagnostidae it possesses numerous peg pits, and in Agnostidae, Ammagnostidae, and Aspidagnostidae it is covered by convex tubercles of different shapes. It is possibly composed from biomineralized plates produced by hypodermal cells. These plates join to each other forming fracture-like polygonal junctions.

The basal part of this layer is composed of a rhomboid-shaped network of fibrils. This network is also visible as an ordered mesh-like texture at the bottom of the pits, where the inner layer becomes thinner. Conversely, it can also be detected as the mesh-like texture of the surface of acrolobes where the inner layer becomes thin.

The basal part with fibrillar mesh can be a protein-chitin matrix produced by the hypodermal cells (Dennell 1976). In one area in the pygidium *Pseudagnostus ex gr. P. cyclopyge*, we found remains of two or three adjusted hypodermal cells with fibrous fabrics on the cells’ surfaces forming from the cell junctions to the middle of the cells (Naimark, Chaika, 2022). The reticular organization of protein-chitinous fibrils is quite common in the cuticle of arthropods (Politi *et al*. 2021). It allows the cuticle to stretch and grow in different directions (a mesh-hammock principle). It is also important that biomineralization of the exoskeleton usually occurs along the protein-chitin matrix (Roer, Dillaman 1984). In this sense, the presence of chitinous fibrils at the base of the cuticle looks quite logical. However, the protein-chitinous matrix of the inner layer has the least chance of being preserved; therefore, it is a rare success to find it in fossil specimens.

In the studied specimens of *P. bulbus, Homagnostus sp. and A. lunulosus*, the inner layer is not as pronounced as in pseudagnostids, primarily due to the absence of multiple peg pits.

In *A. lunulosus,* the layer underlying the outer one could be either transitional or inner. As we can see, this layer in its pygidial spine was composed from separate plates in which biomineralized units went in different directions like in lamellar stacks in the transitional layer of *Pseudagnostus ex gr. P. cyclopyge*. Meanwhile, based on our material we cannot conclude if the inner layer’s plates in *Pseudagnostus ex gr. P. cyclopyge* were also made of differently directed biomineralized units.

In the *Homagnostus sp.* there is a mesh-like texture on the acrolobe and traces of peg pits found on the surface below the outer layer. These features correspond with the inner layer characteristics.

In *P. bulbus,* the layer below the transitional one with the meandering surface is composed of the separate plates that are built of some regular mineralized units. These plates correspond with the plates in the inner layer in *P. ex gr. P. cyclopyge*. In this species the presence of the inner layer seems to be most evident of the non-pseudognostid species.

In the non-pseudagnostid species, there are also pores and various convex tubercles in the inner layer, which are their specific surface feature of the inner layer (Fig. 11B). It needs to be emphasized that their inner layer is thinner and less pronounced than in the pseudagnostid species possibly because of the lighter biomineralization.

The revealed structures of the inner layer, *i.e.* pores of different sizes, the V-shaped groove on the glabellar culmination, peg pits, the basal fibrous mesh, as well as a certain thickness of this layer, could not be imprints or casts of the visceral surface of the outer layer. Thus, these morphological structures indicate the integral identity of the inner layer in the cuticle. The independence of this layer is also evidenced by its elemental composition different in some specimens from the composition in the host rock.

#### Two-layered agnostoid cuticle as previously observed

As shown in the Introduction, it has been previously assumed that the agnostoid cuticle had one layer. However, as we highlighted in our accompanying paper (Naimark, Chaika 2022b) there are still some indications in the literature of an at least two-layered cuticle. In the phosphatized juvenile specimens of Agnostidae (Hong *et al*. 2003, cephalons in figs. 1, 4 and pygidium fig. 10) and in specimens of *A. pisiformis* (Müller, Walossek 1987, pl. 5: 1, 5; pl. 31: 2, pl. 6: 5) the cuticle seems to be splitting at the layers boundary, revealing an underlaying surface different to the upper surface. It possessed multiple convex tubercles lacking in the outer surface. This phenomenon could not be explained in terms of casts and imprints in the rock matrix, in the sense that phosphatized material rock dissolves leaving an undissolved fossilized body, in which original calcite biomineral had been presumably gradually replacing by phosphate (Dalingwater 1991).

Wilmot (1988, 1990) indicated the concave visceral indentations which were not displayed on the outer surface as the specific feature of the cuticle of *H. obesus*. These concavities most probably were the counter-indentations of the convex tubercles of the inner layer, which we observed in *Homagnostus sp.* and *A. lunulosus* and which were evident in the Agnostidae specimens from Korea (Hong *et al*., 2003).

Therefore at least two cuticular layers had been registered in the specimens mentioned in literature but they had been considered to have a single cuticular layer.

Previous studies have focused on revealing prismatic and principal subdivisions in the cuticle similar to those in other modern arthropods (reviewed in Whittington, Wilmot 1997). In agnostids, the cuticle subdivisions apparently do not correspond to these features. The prismatic unit, if present, is extremely thin and limited to the outermost sub-layer with the surface polygons. Also, a pseudo-prismatic appearance may occur if a cuticle is represented only by a transitional layer with stacks of vertical lamellae. Overall, prismatic *versus* principal lamellar layer consideration seems rather senseless adding nothing to understanding how an agnostoid cuticle was built.

### Pores, sensilla, and cuticular glands in agnostids (Fig. 12)

#### <u>Pores</u> (Fig. 12A-H)

Pores are easily recognized from micro-drillings and angular damage in the crystal coating (Miller 1975; McAllister, Brand 1989) by their regular round shape and sometimes by presence of crystal rims.

Pores were found on each of the three cuticular layers. In all specimens, they came in at least two, and sometimes three sizes.

In *Machairagnostus sp.*, the pores were 3–3.5 μm and 1–1.5 μm in size. In *Pseudagnostus sp. ex gr. P. cyclopyge* there were pores of three sizes: 3–5 μm, 1.9–1.5 μm, and 1 μm. 3-5 μm-size pores were found only on the outer layer. The cuticle of *P. josepha* also had pores of three sizes. The smallest pores of 0.5–0.8 μm were located on the inner layer; pores of 1–1.8 μm were located both on the inner and outer layers.

Pores of 1.5–2 µm were recorded in *Pseudagnostus aff. idalis*. *A. lunulosus* possessed pores of 0.5–1 μm and 1.5–2 μm on the outer and inner cuticular layers. In *Homagnostus sp*., in the outer layer, pores of 3 and 7.5 μm were found, while the inner layer contained numerous pores of 1 μm, and funnel-like pores of 7 μm. In *P. bulbus*, relatively large pores (3–4 μm) were located around the median tubercle on the outer layer; the transitional and inner layers had pores of two sizes: 0.5–0.8 and 2-3 μm.

The different sizes of the openings indicate their diverse functions. The pores allow water to penetrate the internal cuticular and hypodermal layers, while conversely allowing mechano- and vibrosensor hairs to pass through them as well (Fornshell 2021).

Pores varied not only in size but also in shape. In *Pseudagnostus sp. ex gr. P. cyclopyge*, in addition to regular pores on the surface layers, two pores were found running parallel to the surface in the outer layer. The walls of these pores, were seemingly built not by calcite crystals, but by chitin, since numerous ordered minute notches are discernible at high magnification in the wall of one of these pores. Such a pattern seems to be a specific display of the pore channels in the protein-chitinous texture of the arthropodial integument.

Some openings on the outer layer are framed by a rim of small crystals; if a pore is located on an acrolobe slope then its crystalline rim might be in the form of a semicircle (Fig. 3J). Conversely, convex smooth ridges instead of crystalline ones were found around the pores on the outer layer. A rim could strengthen the pore walls. The function of such strengthened pore walls is not clear; however we can assume that the reinforced walls of a rimmed pore perform durable transport functions or serve as a passage for mechano-sensory hairs.

An interesting version of pore openings corresponds to the pygidium *P. ampullatus*. Short tubules are arranged in a row between the posterolateral spines along the edge of the border. These tubules form a short and thin marginal ledge. These tubules could be the external shaping of the pores at the edge of the dublura.

Such marginal tubules are known in modern and fossil invertebrates. For example, ostracodes have funnel-shaped radial pore canals in the turndown edge of a dublura. These canals are shaped like tubules going onward, each with a mechanoreceptor hair passing through it (Sohn, Kornicker 1973). Conversely, in the brachiopods Siphonotretidae and Orthida, rows of short tubular spines are located along the shell edge and along its growth lines. It has been supposed that sensorial setae pass through such tubules (Jin *et al*. 2007).

These examples show that tubules could carry sensory elements in agnostids as well. We can also assume that these tubules could connect the interlayer space between the outer/transitional and inner cuticular layers with the external environment. However, it is possible that the tubules were simply protruding edges of the polygonal ridges – that is, they were a side construction product of the dublura turn down. For a more accurate judgment, more observations are required.

In the future, a targeted study of the topography of pores and their thin sections could explain their functions. Our current task is only to confirm the obligatory presence and diversity of these structures on three layers of the cuticle.

#### Sensilla

According to their functions, cuticular sensilla are divided into chemo-, mechano-, and photosensilla, and terrestrial arthropods also have hydro- and thermo-sensors. The sensilla of each functional group are shaped in a certain way, although there is no simple one-to-one correspondence between their shapes and functions.

The shape and function of sensilla in modern arthropods are compared as follows Nowiń Brozek 2019). Mechanoreceptors (including proprioceptors and vibroreceptors) are trichoid, chaetica, styloconic, and campaniform with placoid and digitiform variations. The mechanosensilla, as a rule, are a hollow chitinous bristle or seta sitting in a cuticular indentation; the dendrite passing inside the seta/bristle base reacts to the seta’s/bristle’s bends and deviations from a vertical position. The wall of indentation for the sensillar hairs is built from endocuticle and partially from exocuticle. Campaniform sensilla do not have chitinous outgrowth, but instead are covered from above with a relatively flexible cuticular cap and sub-cup of soft spongy cuticular material. The cap is surrounded by a collar produced of exocuticle and partially from endocuticle. The dendrite passing inside reacts to the change in the shape of the flexible cap.

The chemoreceptor function can be performed by celoconic (coeloconic?), basiconic, and placoid sensilla. They are arranged as a short tubercle sitting in a comparatively deep pit or cavity. The tubercule is rigid, penetrated by one or many pores. Signals from the outside enter an opening/s of the sensilla, to where the dendrite gets. The walls of the sensilla can be single or double; in the latter case, they have a wavy edge in the cross section.

In the cuticle of agnostids, we found structures that have the characteristics of trichoid-chaetica (the length of the setae could not be determined), campaniform and coeloconic sensilla.

There were two examples of trichoid-chaetica sensilla (Fig. 12I–K): in *Machairagnostus sp.* (Fig. 12I) and in *Pseudagnostus aff. idalis* (Fig. 12J). In *Machairagnostus sp.,* this sensillum is expressed as an indentation in the cuticular outer layer, in the center of which there is a very thin wall of the broken tubular structure. This tubular structure is possibly the base of the bristle or chaeta. In *Pseudagnostus aff. idalis*, a hollow broken cone extending from the base of the inner layer is visible. This seems to be the base of the bristle/chaeta on the inner layer, which is supposed to pass through the outer layer that is absent in this specimen.

**Fig. 12.**
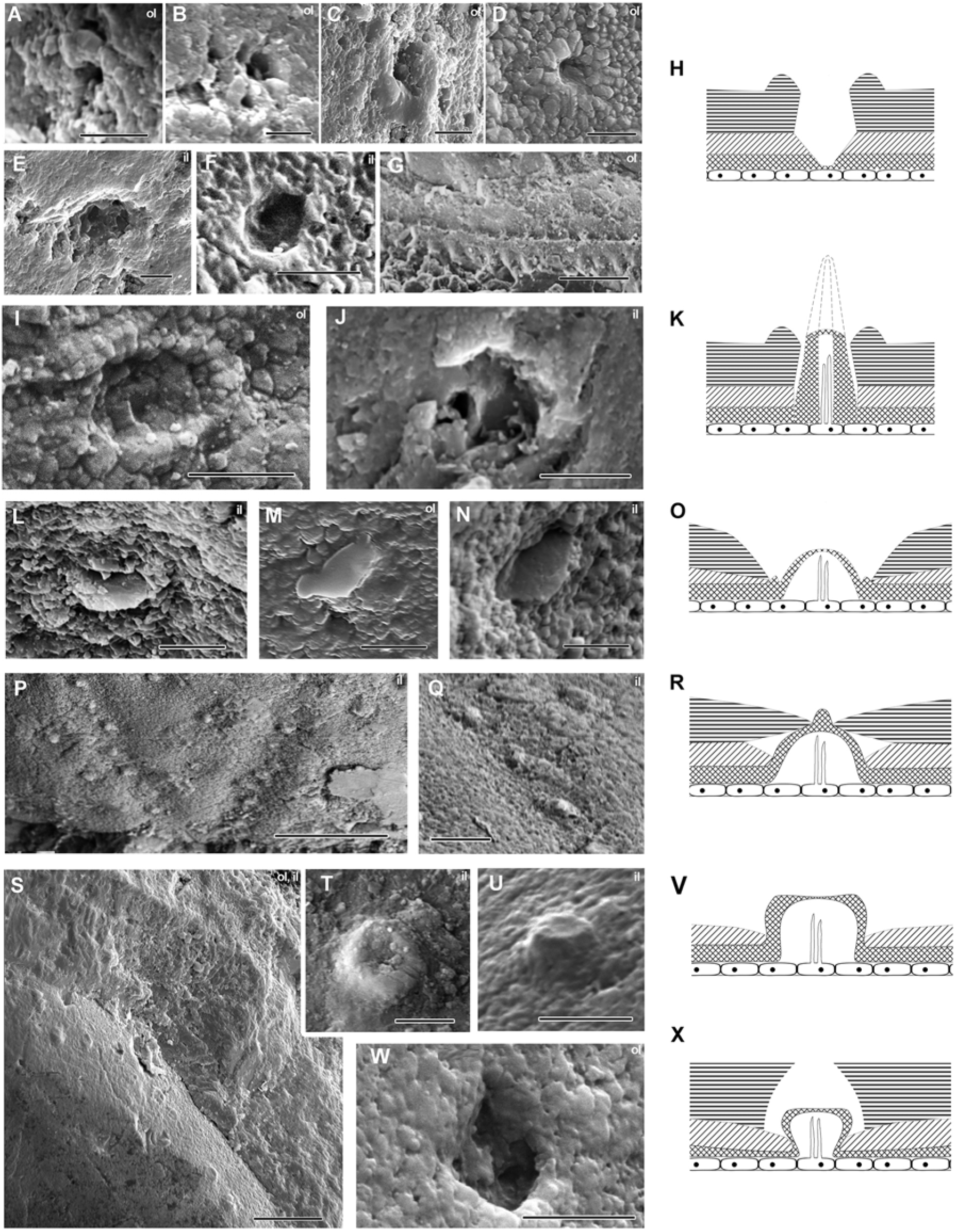
Morphology of cuticular elements in Agnostina. A– D rimmed pores on the outer layer: A, B, *P. josepha*, C, *Homagnostus sp*., D, *Pseudagnostus sp. ex gr. P. cyclopyge*. E, F, openings on the inner layer: E, *Homagnostus sp*., F, *Pseudagnostus sp. ex gr. P. cyclopyge*. G, *P. ampullatus*, tubules on the outer layer between anterlateral spines. H, hypothetical design of a pore going through the outer and inner cuticular layers. I, J, remains of the trichoid/chetica sensilla: I, *Machairagnostus sp*., outer layer; J–*P. aff. idalis,* inner layer. K, hypothetical design of a trichoid/chetica sensillum going through the outer and inner cuticular layers. L–N, campaniform sensilla in *Pseudagnostus sp. ex gr. P. cyclopyge*: L, N – on the inner layer; M – on the outer layer. O, hypothetical design of a campaniform sensillum going through outer and inner cuticular layer. P, Q, V-shape groove with a row of digitiform sensilla in *Pseudagnostus sp. ex gr. P. cyclopyge;* Q, enlarged fragment showing bosses on top of every sensillum dome; only these bosses are visible on the outer layer. R, hypothetical design of a digitiform sensillum going through the outer and inner layers. S, border of the cephalon of *P. josepha*: small elongate tubercles on the inner cuticular surface which sometimes have an indented top. T, V, round convex tubercles with flattened or slightly concave top: T, *Homagnostus sp*.’ cephalic acrolobe; U, in *P. josepha* cephalic border. V, hypothetical design of a round tubercle going through the inner cuticular layer. W, *Machairagnostus sp*.: coeloconic sensillum on the outer layer. X – hypothetical design of a coeloconic sensillum going through the outer and inner cuticular layers. Shadings in the schemes are shown from bottom to top: hypodermal layer with neuronal dendrites, basal fibrous sublayer of the inner layer, principal part of the inner layer, outer layer. Transitional layer is not shown. Scale bars: everywhere 5 μm, except of 50 μm (G, P); 20 μm (S); 10 μm (Q).

Campaniform sensilla are found in abundance on the limb of the pseudagnostid and on the cheeks of *Homagnostus sp*. and *A. lunulosus*. These sensilla are all oval in shape, from slightly elongated to highly elongated in *A.lunulosus*. Some of them are surrounded by a ring at the base (Fig. 12L–N). These sensilla can be deepened in the inner layer of the cuticle (Fig. 12L).

On the inner layer of the cuticular border, there are numerous oval convex tubercles resembling campaniform sensilla in shape (Fig. 12N). They have no expression on the outer layer (and cannot have, since the outer layer is too thick compared to their size), so their function is not clear.

Some of these oval tubercles are irregularly collapsed (Fig. 7H), thus they were seemingly covered with a non-rigid material. In *Homagnostus sp*. with a thin cuticle, smooth elongated bulges are sometimes found on the outer layer, corresponding in size to tubercles on the inner layer. We can only very carefully assume that these bulges are the expression of campaniform sensilla on the outer cuticular layer, and that these elements themselves respond to pressure and/or stretching of the cuticle.

Digitiform sensilla represent a specific, very elongated variation of campaniform sensilla (Fig. 12P–R). In pseudagnostids, they are present in at least the lower part of the V-shape groove and arranged in a series one above the other. Their long axis is parallel to the base of the cephalon. We found V-shaped groove with such sensilla in three samples: in one of them on the outer cuticular layer, and in other two on the inner one (Naimark, Chaika 2022). On the outer layer, only the V-groove itself, very faint parallel traces, and protruding small rounded protuberances are visible, while on the inner layer the elongated sensilla with the rounded tubercles on their tops became expressed.

As a rule, digitiform sensilla function as highly sensitive mechanoreceptors, capturing signals of water currents. Since only the tip of such a sensillum protrudes onto the surface of the cuticle, it probably belongs to some vibroreceptors which could be also equipped with some sensitive hairs. These sensilla are located parallel to the glabellar rear in the groove branches diverging from the center of the base in the glabellar culmination. Such a disposition can indicate a special role for these sensilla: they could register the currents from the symmetrical foramens in the articulating membrane when an animal was closed in the shell. In an open state, this V-groove was possibly covered by the first trunk segment. At least two campaniform sensilla were located close to the base of the V-shape groove along the axis of symmetry (Fig. 12L, M). They probably also belonged to the sensillar group that captured signals of water currents from paired foramens in the articulating membrane.

A special sensory ensemble was found on the zonate border of the pygidium of *A. lunulosus* (see Results). We repeat here that this ensemble included two symmetrical sensory fields on the collar and a series of pores and indentations in the postaxial furrow. It is difficult to imagine any other function for this group, except for the sensory one, although it is impossible so far to guess something more certain concerning its function.

The morphology of the zonate pygidial borders in Aspidagnostidae, as well as in Diplagnostidae, is very specialized and their function is unknown. Given the richness and diversity of the sensory structures of the zonate border in *A. lunulosus*, it is reasonable to assume that the zonation was formed to perform some important sensory functions.

Coeloconic sensilla were found in the outer cuticular layer of *Machairagnostus sp.* They looked like comparatively deep pits with a boss-shaped central element. Сoeloconic sensilla similar in shape with chemosensory functions are known, for example, on antennas in dragonflies (Handique *et al*. 2017).

Coeloconic sensilla are possibly represented by a series of cuticular structures around the glabellar node in *P. bulbus* and axial node in *P. josepha*. Probably, the same sensilla have been mentioned and depicted around the glabellar node on the phosphatized cephalon of *A. pisiformis*, judging by the size and location of the indentations and the presence of internal central elements in them (Müller, Wallossek 1987, p. 38–39, pl. 6:5, 8:6, 8:7, 8:8, 32:5). These structures are arranged in several symmetrically diverging rows going downward from the highest point of the axial node or downward around the glabellar node. On the outer layer, they are visible as indentations, and in the inner layer, they are represented by convex round (*P. bulbus*) or flower-shaped (*P. josepha*) tubercles. In shape, they resemble coeloconic sensilla with a flattened or slightly convex sensory dome; such chemosensory coeloconic sensilla are known, for example, in beetles (Handique *et al*. 2017).

Another kind of numerous cuticular structures were revealed in *Homagnostus sp*. and *P. josepha*. These structures look like a round dome with a central depression or flattened apex (Fig. 12T-V). They are small and located on the inner cuticular layer; no manifestation corresponding to this type was found on the outer layer. Besides, the thickness of the outer layer in *P. josepha* appeared to be 10 times bigger than these sensilla therefore they would have had some very specific exit to the outer surface. There are sensilla – superficially similar to sensilla in Cumacea (Malacostraca) – located on the head along the edge of the rostral region; the function of these sensilla is unknown (Laverack and Barrentios 1985).

We see that various mechano-and chemosensory structures were present on the cuticle of agnostids. The shell was protruded by pores and canals, through which small mechanoreceptor trichae and larger chaetae could pass. The borders were probably a highly sensitive area as mechanosensilla were located in abundance on the borders, and in addition, a series of tubules rimmed the edge of the border between the pygidial posterolateral spines. A special sensitive zone with sensory fields could also be located on the posterior edge of the pygidium. In *A. lunulosus*, this zone is represented by a specialized sensory field on a zonate border and posteroaxial furrow, while in *A. pisiformis*, the sensory function of the of the pygidial rear is emphasized by the accumulation of pores, which are more numerous than elsewhere on the carapace (Müller and Wallossek 1987).

At the glabellar rear, close to the cephalic and first tergite joint, there was a series of sensilla that seemingly detected water currents, for instance from foramens in the articulating membrane. The chemoreceptor zones seemed to be concentrated around the glabellar and axial nodes. Thus, in agnostids, not only the limbs and abdominal side, but also the dorsal side and borders of the carapace were provided with a rich set of sensory structures. As agnosids spent part of their time inside a closed shell, it was important for them to have sensitive structures on the shell surface. Even in the enrolled position, blind agnostids could receive a variety of information about the outside environment.

#### Cuticular glands

Cuticular glands in arthropods are located in the hypoderma, releasing various secretory substances through thin openings in the cuticle. Special hypodermal cells are elongated into tubes that become the walls of the ducts. Inside a duct, there could be a thin rod composed of fibrillar material, along which secretory substances rise up to the surface. If there is no rod, then the secretion simply fills the thin duct.

In the specimen of *P. aff idalis*, we found a structure on the inner layer with the listed features of cuticular glands. It was a fragment of a tube lying near a round recess of the same diameter as the tubular element. A very thin rod is visible in the center of the recess. This rod seems to be either an internal fibrillar structure of the duct or the fossilized secretion itself (Fig. 2L).

We also assigned the peg pits – the most numerous structures of the inner layer of Pseudagnostidae – to glandular formations.

The peg pits are evenly distributed over the surface of the inner layer of the cephalon and pygidium. Only in muscle scar areas and in all furrows were the peg pits few.

The peg pits look as follows. Their basal part is represented by a shallow depression in the chitin-protein fibrillar sublayer, in the center of which there is a tiny bulge. The pit becomes deeper from the surface of the inner layer as its base still reaches the fibrillar sublayer. The peg in the center of the pit may be surrounded by a ring; though on some pegs, the ring was not detected. We assume that there might be some kind of a glandular canal which passed from the fibrous bulge to the opening in the top of a peg. Such a schema can explain the holes in pits where the pegs were destroyed and a tiny (0.1–0.2 μm) opening on the top of one of the investigated pegs. On the surface of the transitional layer, the peg pits were expressed as rounded traces with blurred central structures. On the outer layer, under the polygonal sublayer, the peg pits possibly turned into depressions or became fully effaced.

We tend to attribute the peg pits to glandular rather than sensory structures. There are two reasons for this idea. First, sensilla presenting only on the inner layer of the cuticle are extremely few in arthropods. We know only one such example: dome-shaped chemoreceptors on the endocuticle in mandibulate moth larvae; these sensilla capture chemosignals in the interlayer space (Dupont, 2014). Secondly (and more importantly), there is a gradual smoothing of the pits from the innermost layer to the outermost. A functional sensory structure should have a clear representation on both layers of the cuticle and cannot gradually smooth out from inner layer to the upper surface. Glands, on the other hand, can be smoothed out as the secretion is extracted to the surface. Accordingly, we assume that the peg pits in pseudognostids and possibly in other agnostids are cuticular glands that secrete some substance onto the surface of the cuticle to build the upper layers of the cuticle.

The outer layer in pseudagnostids is much thicker than in the other studied agnostids, thus their cuticular glands are expected to be more numerous and frequently expressed. The tops of the pegs are covered almost everywhere with structureless substance, chemically similar to that of the shell, which could be the biomineralizing materials. The outer and transitional layers were possibly built utilizing the secretion passing from hypoderma to the surface of the inner layer. The outermost sublayer with the polygons possibly formed at the very beginning of the molt, before other cuticular subdivisions: along the cell walls of the hypodermis. This would explain the surface polygonal pattern. In this way we could logically links the available facts, however additional material is needed to confirm and detail shell formation in agnostids.

### Comparison with cuticles in other Arthropoda

The cuticular subdivisions accepted for modern arthropods are epicuticle, procuticle, including exo- and endocuticle, and in chelicerates also mesocuticle. To compare these subdivisions with the agnostoid integument, we first provide brief descriptions of the procuticle arrangement (obviously epicuticle has little chance to become fossilized) in the main classes of arthropods.

In insects, the procuticle consists of exo- and endocuticles with lamellae of different thickness: thinner in the exocuticle and thicker in the endocuticle. Such a ratio of exo- and endocuticle lamellae thickness is characteristic of all modern groups of arthropods. The membraneous layer lies under the procuticle, separating the hypodermis and the intermolting cuticle from the postmolting cuticle. After molting, the membranous layer (ecdysial membrane) acts as a protective outer cover in the forming postmolting cuticle. In insects, in rare cases, the space between the exo- and endocuticle can be filled with fluid. For example, in the mandibulate moth larvae (Micropterigidae) mentioned above (Dupont 2014).

In crustaceans, the constitution of a calcified cuticle resembles those of insects. It is a combination of a thin outer layer (exocuticle) and thick inner layer (endocuticle). The exo- and endocuticle have a lamellar structure: the exocuticle lamellae are always thinner than the endocuticle lamellae (Dillaman *et al*. 2013). Below the exo- and endocuticle lays a non-mineralized membranous layer that separates the cuticular layers from the hypodermis and from the emerging new cuticle of the next instar stages (Roer, Dillaman 2018).

The lamellar organization seems to be uniform in crustaceans, but different mechanisms of lamella biomineralization can induce a variety of cuticular structures. For example, in Xanthidae crabs, the endocuticle is divided into columnar units delimited by organic material. In this case, the boundary between the exo- and endocuticle is highlighted by a specific regular pattern of rounded projections on the surface of the endocuticle (Roer and Dillaman 1984). As a result, the exfoliated surface of the endocuticle looks completely different from the exocuticle: it is a regular pattern of uniform convex tubercles.

In Eurypterida and horseshoe crabs, the shells have a roughly similar organization. In the carapace of horseshoe crabs, the outer layer (exocuticle) is built of horizontal lamellae whose thickness decreases from the upper to the inner layers, becoming thicker again at the exo-/endocuticle interface. The endocuticular layer is relatively thick; it consists of very thin lamellae organized in horizontal, vertical, or slanting units (stacks) that merge into one another (Dalingwater 1973; Dennell 1976; Mutvei 1977, 1981; Ma *et al*., 2022). It is subdivided by an interlayer membrane into two parts with differently organized chitin fibrils. The interlayer membrane itself appears to be dominated by proteins rather than chitin (Dennell, 1976). In some studied specimens of Eurypterida, the inner layer of the cuticle was thinner than the outer one. Mutvei (1977) explained this inverse ratio of the layer thicknesses by dissolving an inner layer in exuvia during molting.

The trilobite integument has been believed to consist of an outer and inner layer, which have sometimes correlated with the exo- and endocuticle of modern arthropods (Dallingwater *et al*. 1999). These layers can be separated by an organic rich interlayer (Dallingwater *et al*. 1991). The exocuticle has been assumed to be composed of prismatic crystals (prismatic layer). However, it was shown for some Cambrian and Silurian trilobites that the outer layer of the cuticle was built from thin lamellae. It is possible that the idea of the prismatic nature of the outer layer in their cuticle has been based on a material with a secondarily recrystallized outer layer or a post-mortem layering of calcite on the initially lamellar structure (Dalingwater *et al*. 1991; 1999).

The trilobite inner layer is comprised of several laminate zones where lamellae differed in thickness. The thinnest lamellae were located in the outermost laminate zone followed by the zone with the thick lamellae, while the inner zone consisted of several thin lamellae (Wilmot 1988, 1990, review in Dallingwater *et al*. 1991). In the phosphatized *Ellipsocephalus polytomus’* cuticle, a thin and high organically rich layer was revealed between the “exocuticle” and “endocuticle” (Dallingwater *et al*. 1991). In the “endocuticle” itself, there were channels 6–7 μm in diameter, which on the surface of the “endocuticle” looked like regular convex rounded tubercles. According to the authors of that study, biomineralizing calcium was excreted through these channels, taking part in the formation of the “exocuticle” (Dallingwater *et al*. 1991; 1999).

Examples of the interspaced cuticle in larvae of mandibulated moths, the cuticle of the trilobite *E. polytomus* with a highly organic interlayer, and the endocuticle of horseshoe crabs separated by an interlayer lamina, suggest that in arthropods the boundaries between cuticular divisions can be accentuated by organic filling and membranes.

The outer and inner cuticular layers of agnostids seemingly have a lamellar structure with lamellae of different thicknesses and directions—the outer lamellae are thicker than the inner ones (at least, in the *P. ex gr. P. cyclopyge* where we were able to investigate them). The lamellae in the outer layer changed their thickness from comparatively wide close to the upper surface to thin close to the transitional layer. The transitional layer consisted of separate stacks of lamellae arranged in different directions (horizontally, vertically, or obliquely). The upper layers and the inner one were interspaced by some kind of rich organic material of unknown nature forming a gap between them.

Thus, the main similarities between the cuticle of agnostids and integuments of other arthropods are the organization of the cuticle from several subdivisions and their lamellar structure, as well as the alternation of zones with thick and thin lamellae. The agnostoid cuticle essentially differs from the cuticle of other arthropods by; (1) the inverse ratio of the thickness of the outer and inner layers; (2) the ready splitting of the cuticle into the upper part and inner layer; (3) the reverse ratio of the thickness of the lamellae in the outer and inner layers: the thicker ones were close to the outer surface top while the thinner ones – to the visceral surface. As can be seen from these brief characteristics, the agnostoid cuticle is most similar to the integument of trilobites, eurypterids, and horseshoe crabs.

Differences in the agnostoid and trilobite integuments are seen in the inverse thickness of thin and thick lamellae: in the trilobite outer cuticular layer (possible exocuticle) the lamellae are thin, while in agnostids the thinner lamellae are located in the transitional layer. However, the trilobite outer cuticular layer included lamellae of variable thickness (thin- and thick lamina zones in Wilmot, 1988). This indicates a high variability in the lamellar architecture of endocuticles in the ancient groups of Arachnomorpha. It is also worth noting that the thickness of the lamellae is associated with the mechanical properties of the cuticle, and it should differ in trilobites and miniature agnostids.

It is worth noting that the trilobite *Hupeidiscus orientalis* (Eodiscida) has two cuticular layers along which the cuticle splits (Li *et al*. 2012) the same way as in agnostids. This feature is not common in arthropods, meanwhile it has been found in miniature agnostids and eodiscids which had been previously encompassed in the group Monomera in Trilobita. Conversely, on the inner layer of *H. orientalis*, there are many small convex tubercles resembling those on the inner layer of the agnostoid, especially on the Agnostidae cuticle. We see in this the certain similarity of the agnostoid and trilobite cuticles.

The agnostoid cuticle also resembles that of the horseshoe crabs and eurypterids in the order of alternation of thin and thick lamellae (thicker ones in the outer layer and thinner in the inner layer). On this feature these groups differ from all other arthropods. Also similar in these groups is the constitution of the transitional layer that is made of stacks of horizontal, vertical or slanting lamellae (Dalingwater, 1975).

If we focus on the structure of the cuticle, the agnostids seem to be closer to some Aracnomorpha group, to trilobites or to Xiphosurans and Eurypterida but not to Crustaceans.

The morphology of the limbs has redirected Agnostina from Trilobita to the stem-group of Arthropoda (Müller and Wallosek 1987). However, at the same time, the tagmatization of agnostids and their “trilobate” shell are similar to those of trilobites, and therefore they were previously united with and derived from eodiscid trilobites. The recent study of agnostids’ limbs suggested that they could still be considered as the descendants of trilobites; the differentiation of agnostids’ limbs could be the result of their high eco-morphological specialization (Moysiuk, Caron 2019). Indeed, if we accept the hypothesis of a trilobite ancestry, then their cuticles will appear to share certain specific features. But still, the agnostoid cuticle looks highly specialized.

The specific structural features of the agnostoid cuticle are possibly associated with their minute size, as well as with a planktonic or near-bottom lifestyle; conversely, adaptation to an in-shell life played a role. Active movement in a water column required a reduction in weight (and consequently in thickness) of the cuticle, while an in-shell existence required certain sensors to receive signals about the outside environment. In agnostids, this kind of thinning affected mostly the transitional and especially inner cuticular layers and the degree of inner layer biomineralization possibly decreased. The evolutionary development of the pseudognostids resulted in a secondary thickening of the shell. However now it was not the inner layer that re-thickened, but the outer one. To form a thick outer layer, the glandular system that delivered mineralizing agents to the outer layer was also improved, and numerous peg pits appeared.

## CONCLUSIONS

The SEM study of even a limited number of specimens has provided rich new material on the integumental anatomy of Agnostina. The layered structure of the cuticle was proved, and a rich ensemble of sensory elements was also revealed. The cuticle of these tiny arthropods seems to be well adapted to their planktonic (swimming) way of life and tuned to receive and convey signals from environment to the body.

We found that the agnostoid cuticle consisted of three layers, of which the inner one was readily split from the upper ones and was possibly interspaced with them by a rich, organic material. The outer layer was relatively thick (15-100 µm), heavily mineralized, and with a pronounced polygonal pattern on its surface. The transitional layer was composed of stacks of think lamellae. The inner layer was built up from thin plates that corresponded to polygonal hypodermal cells; it was possibly lightly mineralized. The inner layer was comprised of two subdivisions; the upper one built upon a mesh-like matrix, and was probably composed of protein-chitin fibrils. According to the lamellae thickness alternation in the cuticular layers, as well as the presence of the stacks of lamellae in the transitional layer, the agnostoid cuticle resembles the cuticle of horseshoe crabs, Eurypterida and trilobites (early Arachnomorpha).

In Pseudagnostidae, the outer layer is the thickest among other studied agnostoid species, and their inner layer bears numerous peg pits. The outer cuticular layer of other agnostids is considerably thinner and peg pits appear as much less pronounced than on the inner layer. The peg pits could be cuticular glands related to the biomineralization of the thick outer layer. They were likely involved in the delivery of substances from the hypodermis to the outer layer.

On the inner layer of non-pseudagnostid cuticles, there are numerous convex tubercules different in size and shape. Their nature is not clear.

The cuticle of agnostids bears a variety of pores and elements that we interpreted as sensory. We found structures which resembled trichoid and campaniform sensilla, for which a mechanosensory function is usually supposed; coeloconic and placoid sensilla with tentative chemosensory properties. On the rear of the cephalons in Pseudagnostidae and in the pygidia of Aspidagnostidae, sensory fields were revealed that have no analogues in other arthropods. The sensory field on the pseudagnostid’s glabellar rear is represented by a series of digitiform vibroreceptors which were possibly associated with some mechanoreceptor tasks given their position in the glabella. The function of the sensory complex on the highly specialized zonate border in Aspidagnostidae is difficult to suppose.

## Acknowledgements

This project would not have been possible without the highly professional help and advices of R.Rakitov who assisted us with the SEM imaging. We also acknowledge the input of the anonymous referees who revised and improved the manuscript.

## DATA ARCHIVING STATEMENT

Data for this study [including x, y and z] are available in the Dryad Digital Repository: https://datadryad.org/stash/share/XXXX

---

**Supplementary Table 1.**
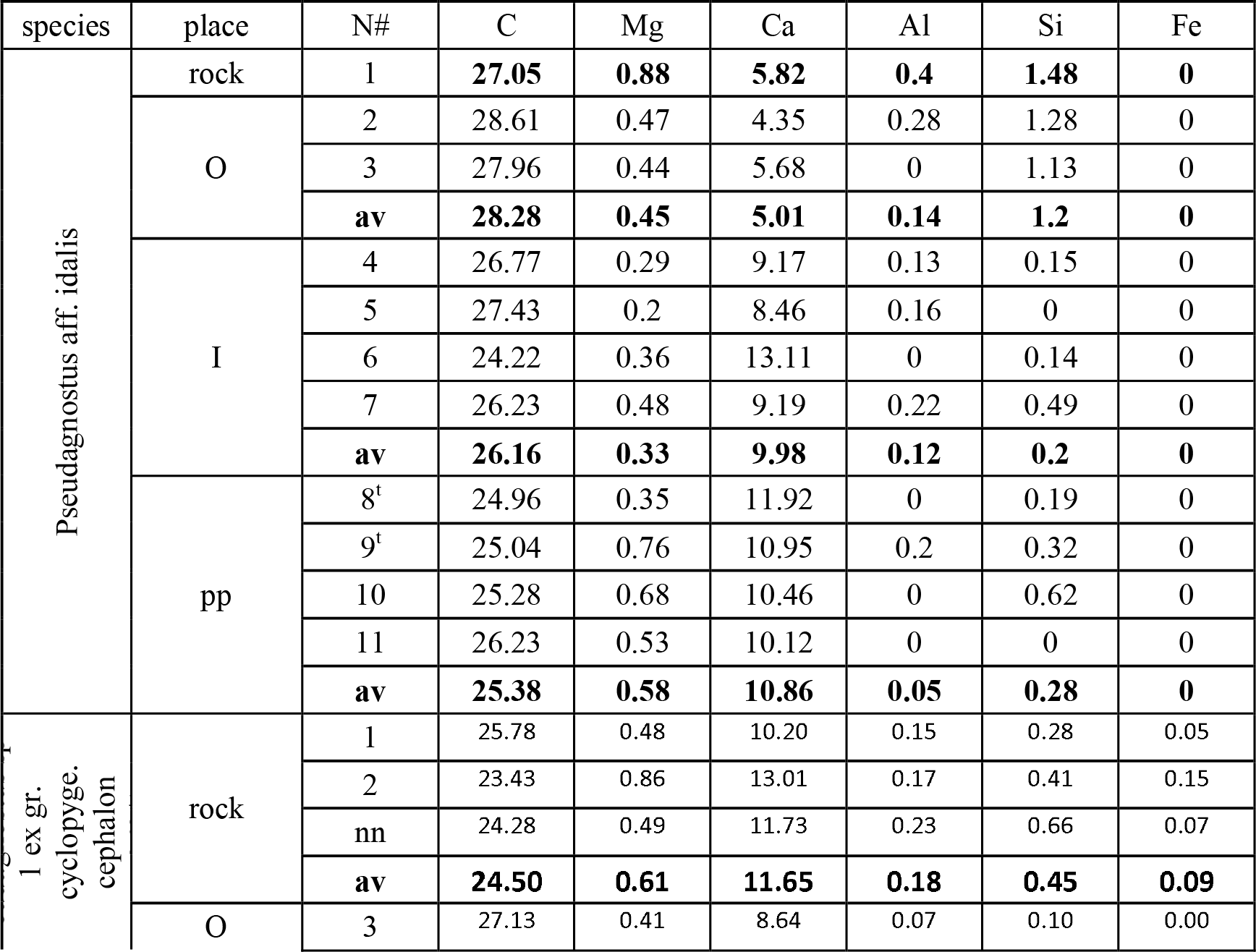
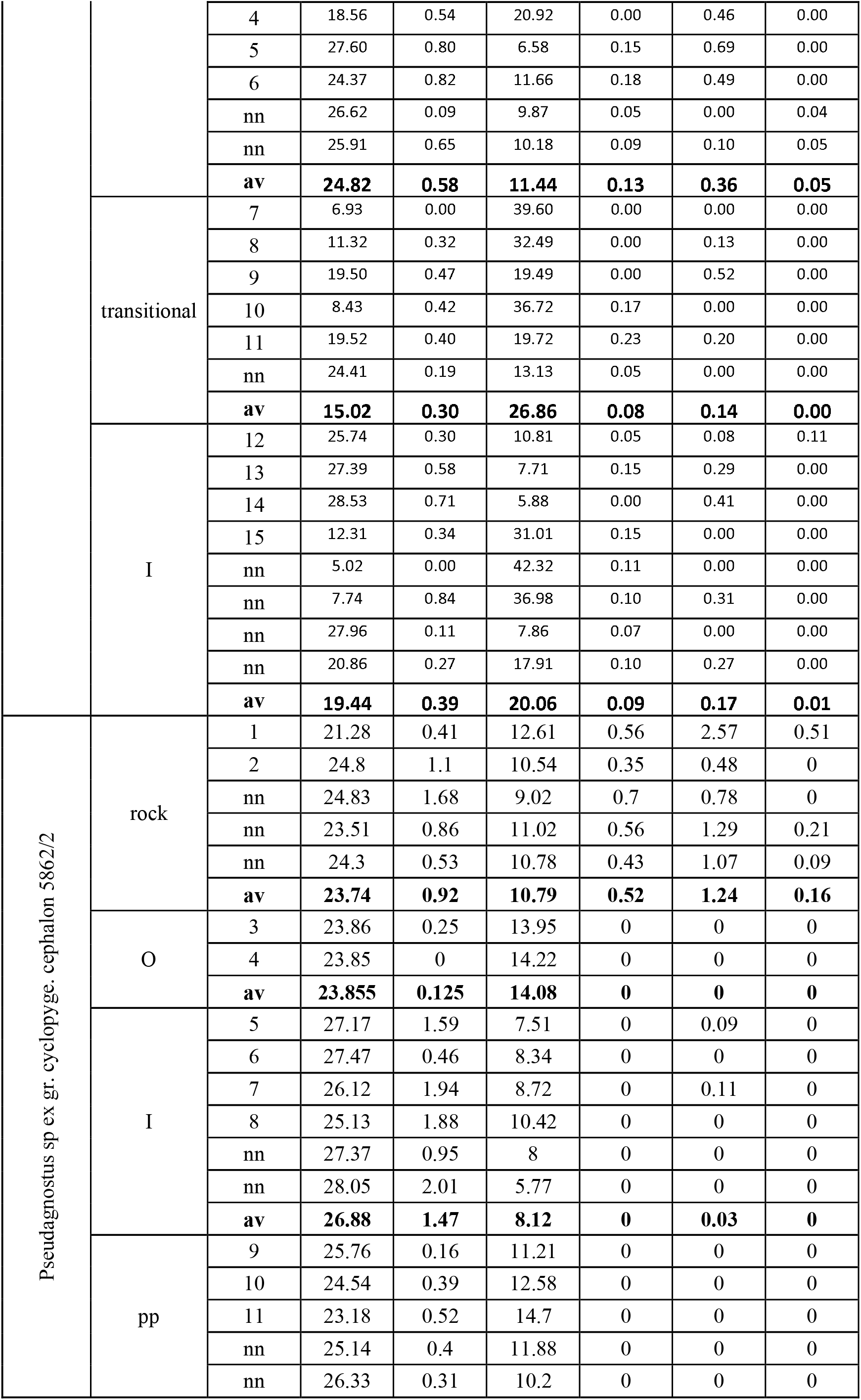
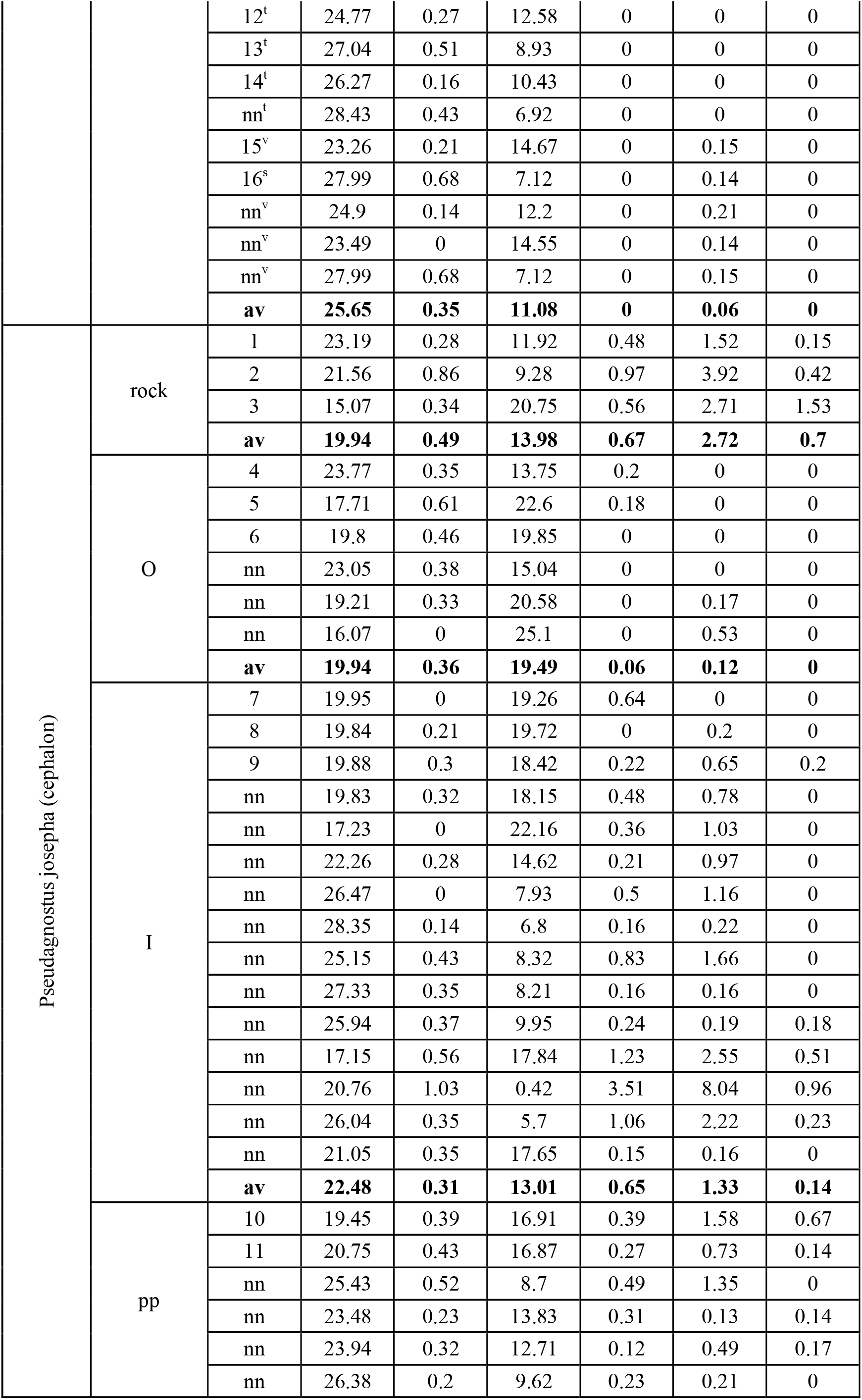
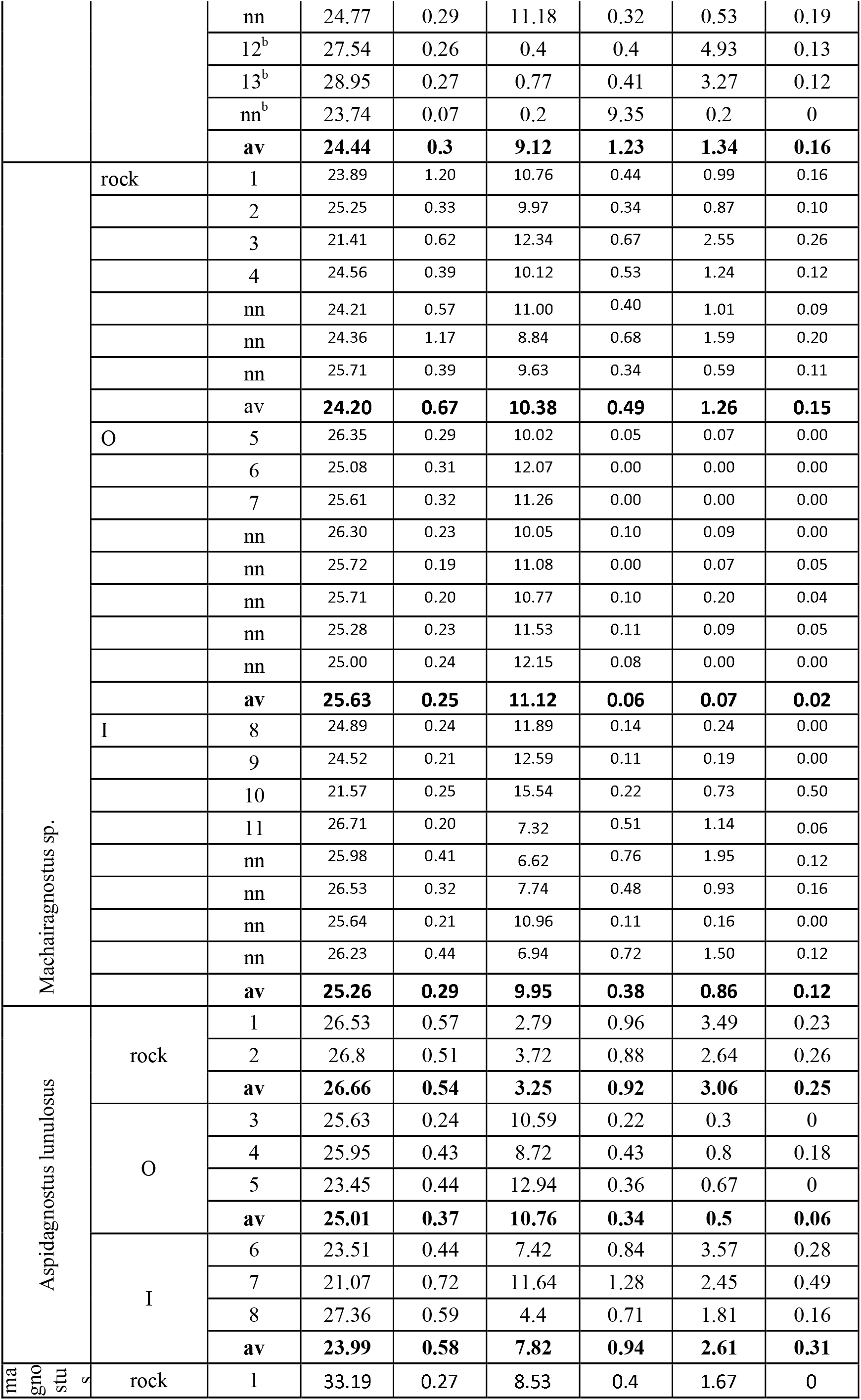
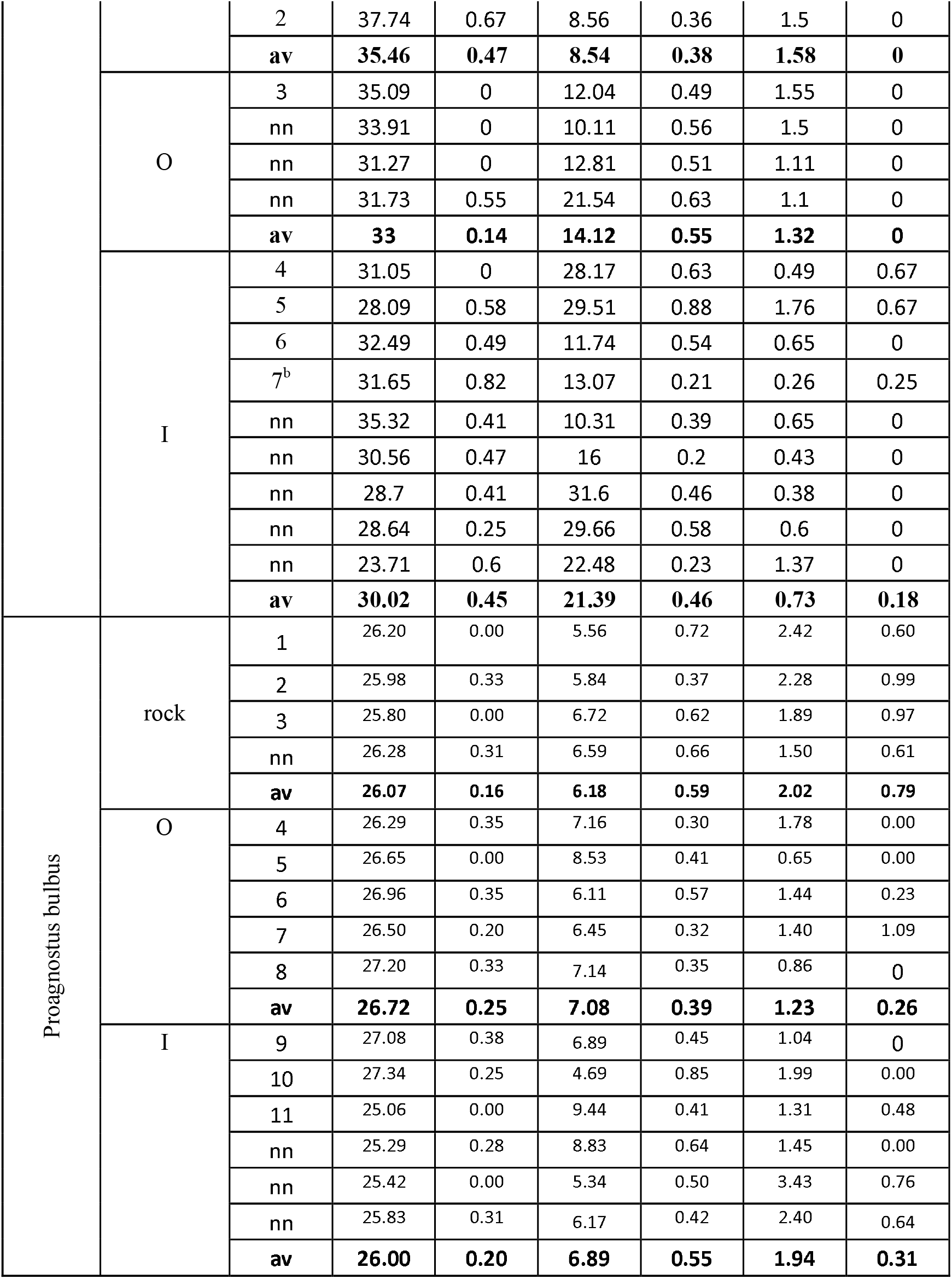
Point elemental analyses of the agnostoid exoskeletons and the host rocks (in atomic %; K, Na are ommited). Measurements for outer (O), inner (I) layers and peg pits (pp) are shown. Numbers correspond to those in Figures 2-10; nn – analyses which were not reflected by asterisks in the Figures; upper indices: t – top whitish matter on the pegs, v-tubercles in the V-shape groove, s – the elongate tubercle between the V-groove branches in the inner layer, b – bulbous structures on the acrolobes.

## Notes

### Competing Interest Statement

The authors have declared no competing interest.

